# Electrode Reduction by *Vibrio natriegens* Depends on Balanced Expression of Multiheme Cytochromes

**DOI:** 10.64898/2026.05.06.723379

**Authors:** Matthew D. Carpenter, Wen-Chia Chen, Caroline M. Ajo-Franklin

## Abstract

Multiheme cytochromes *c* can facilitate electron transfer across the periplasm and outer membrane of Gram-negative bacteria to enable extracellular electron transfer (EET). EET empowers bacteria to maintain redox balance in oxygen-poor environments by donating electrons to solid materials. The resulting electron flux makes EET pathways useful tools for interfacing microorganisms with electronics. Recently, *Vibrio natriegens*, a marine bacterium notable for its rapid growth and expanding biotechnological applications, was found to perform iron reduction in a multiheme cytochrome *c*-dependent manner. However, the role of the *V. natriegens* EET genes in facilitating reduction of electrodes remains unexplored. Through single gene deletion and complementation, we find that each of *cymA*, *pdsA*, *mtrA*, and *mtrB* are required for production of electrical current by *V. natriegens* cultures. Curiously, deletion of the outer membrane decaheme cytochrome *mtrC* diminished but did not abolish electrode reduction. Modulating the induction of expression of *mtrA* and *mtrC* revealed that only a narrow range of induction of these decaheme cytochromes allows balanced cytochrome *c* production and EET. These findings indicate that a multiheme cytochrome-based EET pathway enables *V. natriegens* to reduce electrodes and that this pathway requires carefully balanced gene expression to function. This characterization of the role of multiheme cytochromes in the electroactivity of an emerging microbial chassis for biotechnology will enable new bioelectronic applications for *V. natriegens* and new understanding of the metabolic function of EET.

## Introduction

Microorganisms can reduce extracellular electron acceptors, and a molecular understanding of this process informs our understanding of energy conservation (1) and our ability to harness it for biotechnological applications (2). Extracellular electron transfer (EET) is a metabolic process by which electrons accumulated through cellular catabolism are used to reduce terminal electron acceptors located outside the cell envelope (3). EET can enhance cellular regeneration of oxidized electron carriers, sustaining cellular metabolism under conditions where cell-permeable terminal electron acceptors are unavailable (4, 5). While the first microorganisms found to perform EET (e.g., *Geobacter* and *Shewanella* species) use it as a primary mode of conserving energy (4–6), there is an increasing recognition that some microorganisms use EET as a secondary (7–9) or latent (10) energy metabolism under specific environmental conditions. As more microorganisms inhabiting varied ecological contexts are found to utilize diverse EET, understanding the relationship between the metabolic functions of EET and the structures that compose EET pathways becomes increasingly important (11). Pathways that evolved to perform EET to environmentally relevant electron acceptors, such as solid iron oxides, can often perform EET to polarized electrodes, generating electrical current (12). Consequently, biological engineering of electroactive microbes can produce bioelectronic systems for environmental sensing (13), bioelectrocatalysis (14), bioremediation (15), power generation (16), and synthesis of polymeric materials (17).

EET activity has recently been observed in *V. natriegens* (18, 19), presenting intriguing questions about this organism’s metabolic strategies. *V. natriegens* possesses fermentation pathways for anaerobic growth (20), can grow extraordinarily quickly (under ideal conditions, its doubling time is less than 10 minutes) (21), and is believed to have a highly reductive metabolism (22), suggesting the metabolic role of EET may differ from that observed in model EET-performing organisms. Furthermore, Gemünde, *et al.*, observed electrical current production from *V. natriegens* supplied with glucose and a positively polarized electrode (23), offering opportunities for the development of microbial bioelectronic systems using *V. natriegens*. This potential is notable because *V. natriegens* is also a versatile emerging chassis for biotechnology (24), with applications in biodegradation (25) and bioproduction (20, 26, 27). Understanding the relationship between EET and anaerobic metabolism in *V. natriegens* may unlock applications for *V. natriegens* in bioelectrocatalysis.

*V. natriegens* possesses genes that support EET to iron (18). Conley, *et al.*, described the ability of *V. natriegens* to perform EET to ferric iron and found that *cymA*, *pdsA*, and the *mtrCAB* operon were required for iron reduction (18). These genes have substantial homology to genes within a common archetype for Gram-negative EET pathways – the use of a series of multiheme cytochromes *c* and a porin-cytochrome *c* complex to move electrons from inner membrane quinols to the exterior face of the outer membrane (18, 28, 29). The gene products of *mtrCAB* of *V. natriegens* are homologous to those of the *mtrCAB* operon of *S. oneidensis*, which encodes an outer membrane-localized complex in which a β-barrel porin (MtrB) facilitates interaction between the periplasmically-localized decaheme cytochrome MtrA and the extracellularly-localized decaheme cytochrome MtrC (18). To deliver electrons to MtrA, *S. oneidensis* relies upon an inner membrane-tethered tetraheme cytochrome (CymA), to which *V. natreigens* CymA shares homology, to oxidize inner membrane quinols and reduce one or more soluble periplasmic shuttles that ferry electrons on to MtrA (18). However, the *pdsA* gene product does not share homology with either of the periplasmic cytochrome shuttles of *S. oneidensis*; *V. natriegens* PdsA is homologous to the periplasmic diheme cytochrome shuttle PdsA from *Aeromonas hydrophila* (18, 30). In this way, the set of EET components found in *V. natriegens* represents an interesting “hybrid” of two groups of Mtr pathways (28). Notably, the EET genes of *V. natriegens* are found without accompanying homologues to some of the other EET genes (ex. *mtrDEF*) found in the *S. oneidensis* genome (31). The set of cytochromes necessary to perform EET not only varies between organisms, but also between terminal electron acceptors. Studies in *S. oneidensis* have found that the importance of the outer membrane cytochrome MtrC to EET can vary for different terminal electron acceptors (12, 32, 33). Accordingly, we sought to determine which genes are involved in EET to electrodes in *V. natriegens*, an inquiry that could enable novel bioelectronic applications, such as in biosensing (13).

The observation that the archetypal Mtr pathway can be found in bacteria with diverse ecological niches and metabolic capabilities (11) raises the question of how the regulation of the multiheme cytochromes differs across contexts. Cells that encounter oxygen can face risks from the expression of redox proteins. Dysregulation of cytochrome *c* maturation risks releasing free heme, enabling Fenton chemistry to create reactive oxygen species (ROS) (34); the many hemes of MtrA and MtrC potentially render production of these proteins even more threatening to the cell in the presence of oxygen. Unlike in *Escherichia coli*, the *V. natriegens* cytochrome *c* maturation machinery is not known to be controlled by the oxygen sensor FNR (35, 36). Cytochromes *c* can therefore be produced under aerobic conditions (36), suggesting that *V. natriegens* may rely on other mechanisms to regulate multiheme cytochrome production or prevent ROS.

Regulation of multiheme cytochrome *c* production is also influenced by the need for post-translational modification. Processing by the cytochrome *c* maturation pathway can serve as a bottleneck on the expression of holo-cytochromes *c*. Efforts to heterologously express the *S. oneidensis* porin-cytochrome *c* complex in other chassis have found that increasing transcriptional induction can diminish both EET activity and cytochrome expression (37). Studies in *E. coli* have also found that expression of these cytochromes is limited by the activity of the native cytochrome *c* maturation machinery (38). Similar constraints might be found in *V. natriegens*, a possibility suggested by the report, from Conley, *et al.*, that attempts to express *mtrCAB* from a plasmid within a Δ*mtrCAB* background were unsuccessful (18). Therefore, we sought to investigate what expression regimes of multiheme cytochromes are permissive for functional EET. Elucidation of the multiheme cytochrome balance that supports electrode reduction is critical to enable successful engineering of the EET pathway of *V. natriegens* for bioelectronic applications.

Here, we interrogate the hypothesis that the balanced expression of each of the genes *cymA*, *pdsA*, *mtrA*, *mtrB*, and *mtrC* of *V. natriegens* enables electrode reduction. To assess the contribution of each of the five EET genes to electrode reduction, we individually delete and complement the expression of each gene and measure current production in a bioelectrochemical system (BES). Then, we investigate the limitations on the expression of the cytochromes within the MtrCAB complex by modulating transcriptional induction of MtrA and MtrC and characterizing the effects on EET activity and the expression of other EET genes. These data provide insight into the set of genes and the balance of their expression required for electrode reduction in *V. natriegens*, revealing a need for careful balance of decaheme cytochrome *c* expression.

## Results

### *V. natriegens* expresses Mtr cytochromes and can reduce an electrode

The *cymA* gene (predicted to encode an inner membrane-localized tetraheme cytochrome *c*), *pdsA* gene (predicted to encode a periplasmically-localized diheme cytochrome *c*), and *mtrCAB* operon (predicted to encode an outer membrane-localized porin:cytochrome *c* complex) were previously shown to be required for *V. natriegens* to reduce Fe^3+^ **(Figure 1A)** (18). To facilitate study of this EET pathway, we sought to culture *V. natriegens* in a medium that supports robust expression of these multiheme cytochromes. Fuchs, *et al.*, recently observed that culturing *V. natriegens* aerobically in rich ZYM-5052-v2 medium supported expression of *c*-type cytochromes (36). To test this culturing method, we grew wild-type (WT) and Δ*mtrCAB V. natriegens* using the method of Fuchs, *et al.*, performed SDS-PAGE on whole-cell lysates, and visualized all proteins containing covalently bound heme through enhanced chemiluminescence (ECL). WT cells produced two bands (with approximate apparent molecular weights of 40 kDa and 90 kDa) that were absent from the lysates of cells lacking the *mtrCAB* operon **(Supplemental Figure 1)**. We attribute the higher molecular weight band to MtrC (predicted MW: 87.828 kDa) and the lower molecular weight band to MtrA (predicted MW: 40.038 kDa). The presence of clearly visualizable MtrA and MtrC bands in the lysate of WT cells demonstrates that aerobic culturing in ZYM-5052-v2 supports expression of the decaheme cytochromes required for EET.

**Figure 1:**
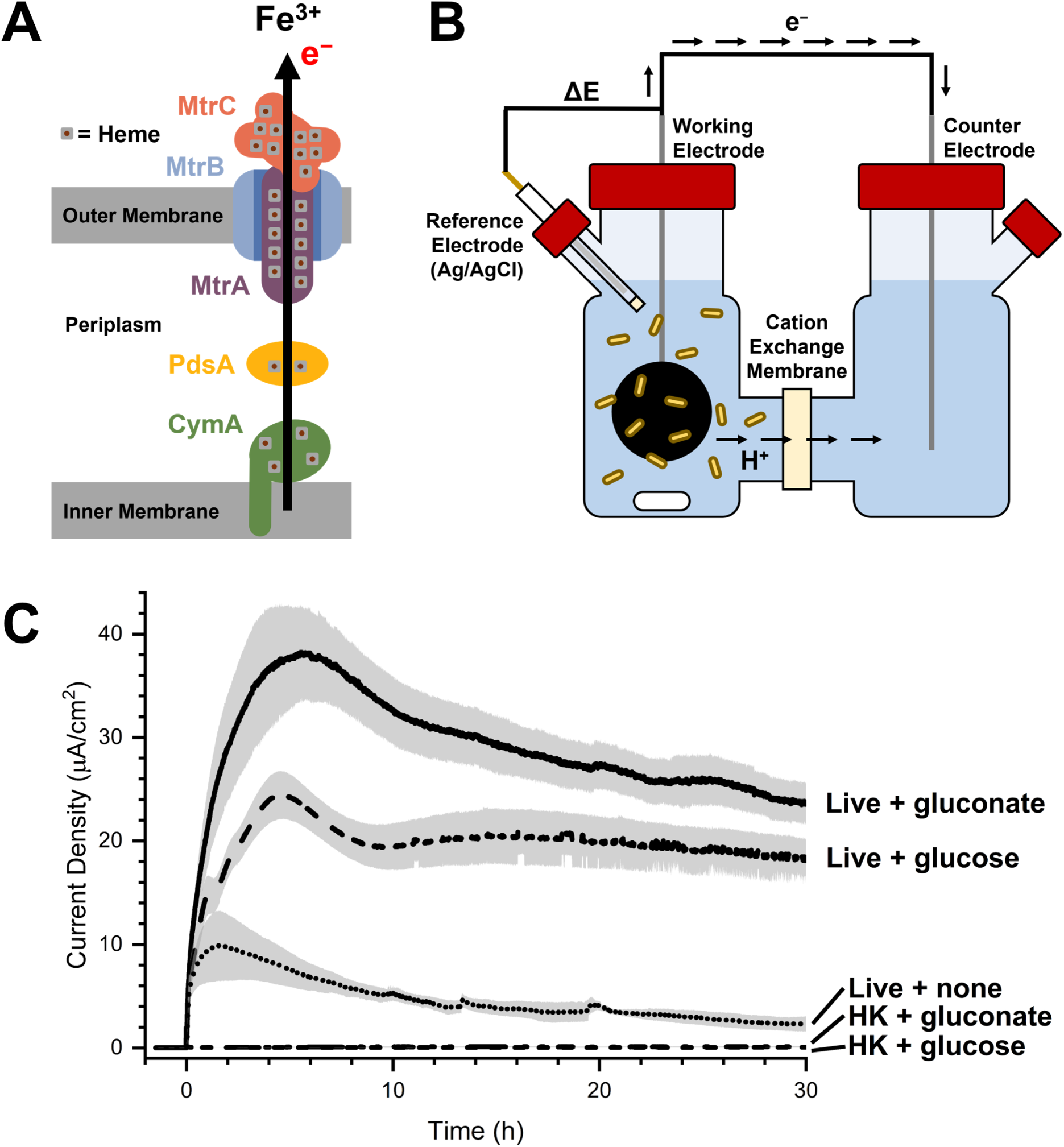
*V. natriegens* produces current in a bioelectrochemical system with either glucose or gluconate as an electron donor. **(A)** A five-gene EET pathway in *V. natriegens* supports ferric iron reduction. The inner membrane tetraheme cytochrome *c* CymA is expected to oxidize inner membrane quinols and pass electrons to the periplasmic diheme cytochrome *c* PdsA, which will in turn reduce the periplasmic decaheme cytochrome *c* MtrA. Electrons are predicted to cross the outer membrane by passing from MtrA to the extracellular decaheme cytochrome *c* MtrC, which contacts MtrA through the outer membrane porin MtrB. **(B)** Electrode reduction was measured using a two-chamber, three-electrode BES in which a carbon felt working electrode was maintained at +200 mV vs. a Ag/AgCl reference electrode using a potentiostat. **(C)** Chronoamperometry of WT *V. natriegens* shows that live cells supplied with 55 mM gluconate (solid line) or 55 mM glucose (dashed line) produce greater current than live cells without electron donor (dotted line) or heat-killed cells with either gluconate (dashed & dotted line) or glucose (short dashed line). Each curve represents the mean of three independent biological replicates. Error bars indicate standard error of the mean.

To sustain current production in a BES, electroactive microbes require a chemical electron donor. Gemünde, *et al.*, observed electrode reduction by *V. natriegens* supplied with ∼55 mM glucose (23). By contrast, Conley, *et al.*, reported that 10 mM glucose did not support significant levels of iron reduction, and identified 10 mM gluconate as an electron donor supporting robust EET (18). To confirm electrode reduction and identify a suitable electron donor for electrode reduction, we measured the electrode reduction of living and heat-killed *V. natriegens* in the presence of 55 mM glucose or gluconate using a BES **(Figure 1B)**. Virtually no current was observed from heat-killed cells supplied with either electron donor. By contrast, the current produced by live cells supplied with either electron donor increased rapidly for 5-6 hours after cells were added to the BES **(Figure 1C)**. Cells with gluconate reached a maximum current density of 38 µA/cm^2^, while those with glucose produced a maximum current density of 24 µA/cm^2^. These currents remained fairly stable with very gradual decline for another 24 hours before the measurements were halted. Live cells without gluconate or glucose showed greater electrode reduction than dead cells, but the observed maximum current density of 10 µA/cm^2^ was far lower than that produced by cells provided with an electron donor. We therefore conclude that electrode reduction by *V. natreigens* is dependent upon cellular catabolism and is supported by both gluconate and glucose. However, gluconate generates higher current and was used for all subsequent experiments.

### CymA and PdsA are required for electrode reduction, but do not affect growth

To investigate the contribution of each putative EET gene to electrode reduction, we designed and constructed single-gene deletion and complementation strains for each of *cymA*, *pdsA*, *mtrA*, *mtrB*, and *mtrC*. All single-gene deletions were scarless deletions. When seeking to restore expression of an EET gene into its deletion strain, we sought to achieve stable maintenance of the gene without the need for selective markers by integrating a synthetic gene expression cassette into a chromosome. We performed chromosomal integration within a 1,500 bp intergenic region on chromosome 2, between genes PN96_RS17100 and PN96_RS17105 (this insertion locus is hereafter referred to as LP1) to minimize the likelihood of disrupting cell function. To further minimize disruption, we flanked the gene expression cassette with strong, bidirectional double terminators (39). To provide strong constitutive expression of the EET gene of interest, we used the P_cymRC_ promoter (without expressing its cognate repressor, *cymR*) (40) and a strong RBS (41). All deletion and complementation strains were sequence verified by sequencing PCR amplicons of the modified locus.

In *V. natriegens*, *cymA* and *pdsA* are homologous to the inner membrane EET gene of *S. oneidensis* and the periplasmic EET gene of *A. hydrophila*, respectively. To evaluate the role of *cymA* and *pdsA* in electrode reduction, we performed chronoamperometry in BESs, comparing single deletion of *cymA* (Δ*cymA*) with its complementation (*cymA*^+^) and single deletion of *pdsA* (Δ*pdsA*) with its complementation (*pdsA*^+^), using live WT and heat-killed WT cells as controls. As observed previously, addition of live WT cells to the BES produced a strong and immediate current response that peaked about six hours after addition. In contrast, heat-killed WT cells produced no detectable response, as expected **(Figure 2A, B)**. The Δ*cymA* mutant showed a significant reduction in current density compared to WT and had minimal residual EET activity. In contrast, the current production of the *cymA*^+^ strain was restored to WT levels and exhibited a comparable temporal pattern that included a rapid onset of current, a peak magnitude similar to the WT, and maintenance of a stable plateau **(Figure 2A)**. Similar trends were observed for *pdsA*. The Δ*pdsA* mutant exhibited reduced current density with no detectable peak, while the current production of the *pdsA*⁺ strain approached WT levels and displayed a similar temporal profile with a rapid initial increase that transitioned to a sustained stable plateau **(Figure 2B)**.

**Figure 2:**
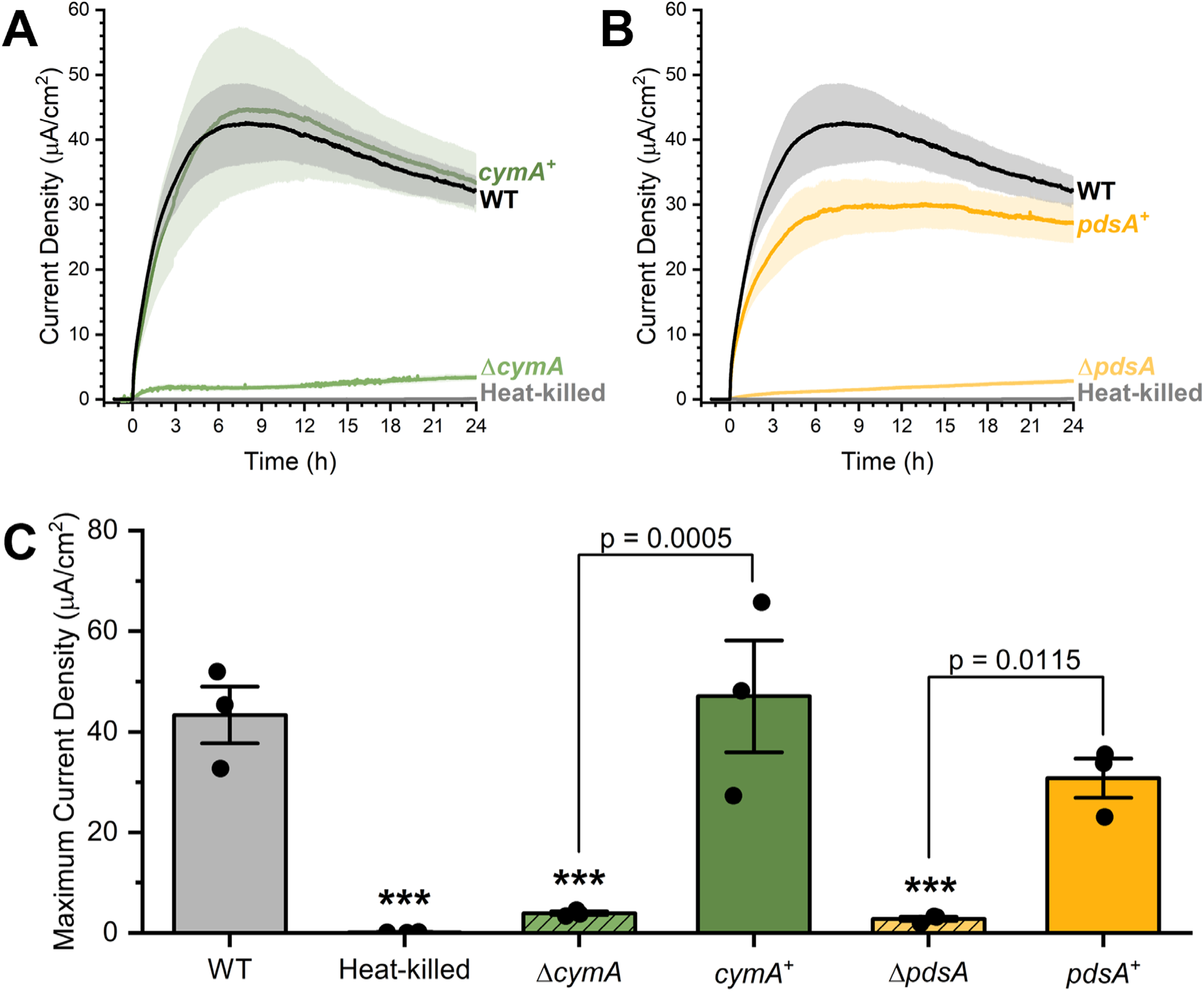
Electrode reduction depends on CymA and PdsA. Chronoamperometry of live (black) and heat-killed (gray) WT cells compared to **(A)** Δ*cymA* (light green) and *cymA^+^* (dark green), or **(B)** Δ*pdsA* (light gold) and *pdsA^+^* (dark gold) cells. Each curve represents the mean of three independent biological replicates. **(C)** Maximum current densities from the time courses reported in **A, B**, demonstrating that removal of *cymA* or *pdsA* largely eliminates current production, and complementation restores current production to levels similar to WT. P-values are calculated by Tukey HSD test. *** denotes p < 0.001 when compared to WT. Error bars indicate standard error of the mean.

We also analyzed the maximum current density to quantitatively compare the magnitude of EET between strains. As expected, live WT cells exhibited a strong maximum current density (43.34 μA/cm²) while heat-killed WT cells showed an extremely low current density (0.20 μA/cm²) **(Figure 2C)**. The Δ*cymA* and Δ*pdsA* mutants exhibited minimum EET, producing maximum current densities of 3.93 μA/cm² and 2.90 μA/cm², respectively. In contrast, complementation with *cymA*^+^ or *pdsA*^+^ restored current production, reaching 47.06 μA/cm² and 30.80 μA/cm², respectively. When compared to their respective deletion strains, both *cymA*⁺ and *pdsA*⁺ complementation strains exhibited a significant recovery in current production (p = 0.0005 and p = 0.0115, respectively) **(Figure 2C)**. The results demonstrate that the gene products of *cymA* and *pdsA* are responsible for the observed electrode reduction phenotypes.

The ability to harness solid extracellular electron acceptors in the absence of oxygen may provide a growth advantage to *V. natriegens*. Conley, *et al.*, observed that EET-capable cells had improved survival 3-6 days after inoculation under iron-reducing conditions when compared to cells lacking genes for iron reduction (18). To assess the effect of EET genotype on the dynamics of growth with electrodes, we cultured Δ*cymA, cymA*^+^, Δ*pdsA*, *pdsA*^+^, WT, and heat-killed WT cells in BESs and sampled the medium every three hours to measure OD_600_. BESs inoculated with live cells showed an increase in biomass over 24 hours of operation, regardless of *cymA* or *pdsA* genotype **(Supplemental Figure 2)**. Over this period of time, typical features of microbial growth curves were observed from all strains. OD_600_ did not increase for a lag phase of multiple hours before the OD_600_ rose rapidly in the window of the 9-, 12-, and 15-hour observations. By the end of the 24-hour experiments, OD_600_ ceased to increase, indicating stationary phase. To ascertain whether any of the tested genotypes demonstrated any differences in growth dynamics, key growth curve parameters were extracted from microbial growth models fitted to the OD_600_ data **(Supplemental Figure 2)**. No significant differences were found among the growth rate **(Supplemental Figure 3A)**, duration of lag phase **(Supplemental Figure 3B)**, or OD_600_ in stationary phase **(Supplemental Figure 3C)** parameters of the strains. The minimally electroactive Δ*cymA* and Δ*pdsA* strains nonetheless grew under our BES conditions, demonstrating that EET activity does not significantly promote growth. We anticipate that NAD^+^ regeneration in the absence of EET is facilitated by fermentation. Our results support a description in which EET does not confer improved anaerobic growth on gluconate.

We hypothesized that CymA and PdsA are mutually interdependent within the *V. natriegens* EET pathway, predicting that a double knock-out would not yield additional losses in electrode reduction capability. To test this, we assessed electrode reduction activity among Δ*cymA*, Δ*pdsA*, and Δ*cymApdsA* strains. Consistent with previous findings, Δ*cymA* and Δ*pdsA* cells showed a stable plateau in low electrode reduction activity (lower than 5 μA/cm²) **(Supplemental Figure 4)**. Injection of Δ*cymApdsA* cells to the BES produced similarly low current density. The results suggest that the contribution of *cymA* to electrode reduction requires *pdsA*, and the contribution of *pdsA* to electrode reduction requires *cymA*. The consistent residual electrode reduction observed in the absence of CymA or PdsA appears to be caused by a mechanism independent of the known EET pathway of *V. natriegens*.

### The outer membrane porin MtrB is required for electrode reduction

Previous investigation of iron reduction by *V. natriegens* examined the effect of deleting the full *mtrCAB* operon (18). Here, we sought to investigate the contribution of each of the genes in the outer membrane complex, the decaheme cytochromes MtrA and MtrC and the beta-barrel porin MtrB, to electrode reduction. As before, we generated mutants of *V. natriegens* with a single gene deletion of *mtrB* (Δ*mtrB*) and with the *mtrB* gene restored under strong constitutive expression at LP1 (*mtrB*^+^) and measured their electrode reduction. In the absence of *mtrB*, *V. natriegen*s produced far less current (3.65 μA/cm²) than was observed from wild-type (WT) cells (24.65 μA/cm²) **(Figure 3A, B)**. When the *mtrB* gene was complemented, current density with similar magnitude (26.77 μA/cm²) and dynamics to WT was produced **(Figure 3A,D)**. These results reveal that electrode reduction heavily depends on MtrB. The recovery of the WT electrode reduction phenotype by complementation demonstrates that the electrode reduction defect of *mtrB* deletion is caused by the absence of the *mtrB* gene product, not by disruption of the rest of the *mtrCAB* operon sequence.

**Figure 3:**
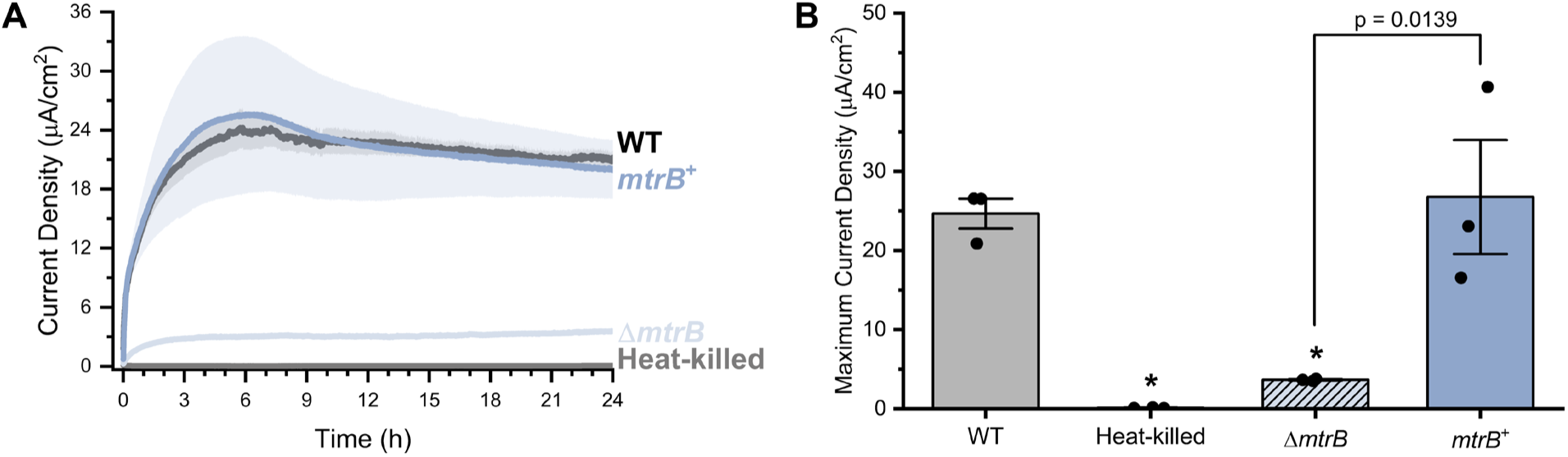
Electrode reduction depends on MtrB. **(A)** Chronoamperometry of live (black) and heat-killed (gray) WT cells compared to Δ*mtrB* (light blue) and *mtrB^+^* (dark blue) cells. Each curve represents the mean of three independent biological replicates. **(B)** Maximum current densities from the time course reported in **A**, demonstrating that removal of *mtrB* largely eliminates current production, and complementation restores current production to levels similar to WT. P-values are calculated by Tukey HSD test. * denotes p < 0.05 when compared to WT. Error bars indicate standard error of the mean.

### MtrA and MtrC contribute to electrode reduction when moderately expressed

To examine the individual contributions of the decaheme cytochromes to EET, we generated single gene deletions for both *mtrA* and *mtrC* and complemented each of these genes in their respective deletion strains, using the same genetic design as was successfully employed for *cymA*, *pdsA*, and *mtrB*. However, both of these complementation strains displayed high instability. We were unable to propagate these strains in liquid culture without losing decaheme cytochrome expression to mutation. Our observations appear to align with the experience of Conley, *et al.*, who reported that “attempts to complement [Δ*mtrCAB*]… were unsuccessful” (18). Producing multiheme cytochromes *c* requires substantial post-translational processing, including secretion, maturation, and complex formation (42, 43). Dysregulation of the expression of cytochromes *c* can impair cell growth (37). We hypothesized that the high level of expression supported by the unrepressed P_cymRC_ promoter rendered complemented expression of *mtrA* or *mtrC* too burdensome for *V. natriegens* to stably maintain. Therefore, we re-designed the expression cassette to enable lower expression levels and be more resilient to evolutionary instability.

To moderate the expression level of *mtrA* and *mtrC*, we expressed CymR, the cognate repressor for P_cymRC_ (40), via genome integration. The *cymR* gene was integrated, with flanking double terminators, within a 729 bp intergenic region on chromosome 1, between genes PN96_RS06225 and PN96_RS06230 (this insertion locus is hereafter referred to as LP2) (44). Expression was controlled by a medium-strength promoter-RBS combination (45). As an additional precaution against evolutionary instability, we swapped the P_cymRC_ promoter sequence for a variant less likely to undergo recombination (P_cymRD_) (46). Using this improved design, we were able to construct and propagate strains with sequence-validated gene expression cassettes for *mtrA* or *mtrC* integrated at LP1. Given that constitutive expression of *cymR* was required to build the *mtrA* and *mtrC* complementation strains, we modified the design of the Δ*mtrA* and Δ*mtrC* strains to also include the chromosomal integration of the gene expression cassette for *cymR* and of the regulatory elements that were used to control *mtrA* or *mtrC* gene expression (these strains are hereafter referred to as Δ*mtrA*-Empty Cassette and Δ*mtrC*-Empty Cassette, respectively).

We then evaluated the electrode reduction ability of this stable suite of Δ*mtrA*-Empty Cassette, *mtrA*^+^, Δ*mtrC*-Empty Cassette and *mtrC*^+^ cultures, alongside WT cells and samples of inactive heat-killed cells. Because we had observed that expressing either of the decaheme cytochromes from an unrepressed promoter resulted in evolutionary instability, we hypothesized that the expression of the decaheme cytochromes might require careful balancing. Therefore, we induced the *mtrA*^+^ and *mtrC*^+^ strains with a range of cumate concentrations and measured the electrode reduction and cytochrome *c* abundance of the cells. While WT cells produced a maximum current density of 31 μA/cm², cells lacking *mtrA* produced current much lower than WT (max. 2 μA/cm²) **(Figure 4A)**. Low concentrations (0 µM and 1 µM) of cumate produced the greatest current density **(Figure 4A)**. This current was greater than that produced by the Δ*mtrA*-Empty Cassette strain but was only around 60% that of WT cells. These results indicate that MtrA contributes to electrode reduction. They also indicate that some basal expression of MtrA occurs in the absence of cumate. Surprisingly, induction with higher concentrations of cumate (10 µM and 100 µM) led to very low levels of current, similar to those observed from cells entirely lacking a gene essential for EET. Despite the lack of electrode reduction, the cells treated with 10 µM or 100 µM cumate showed a prominent band at the expected MW for MtrA when their cytochromes *c* were visualized **(Supplemental Figure 5A-C)**.

**Figure 4:**
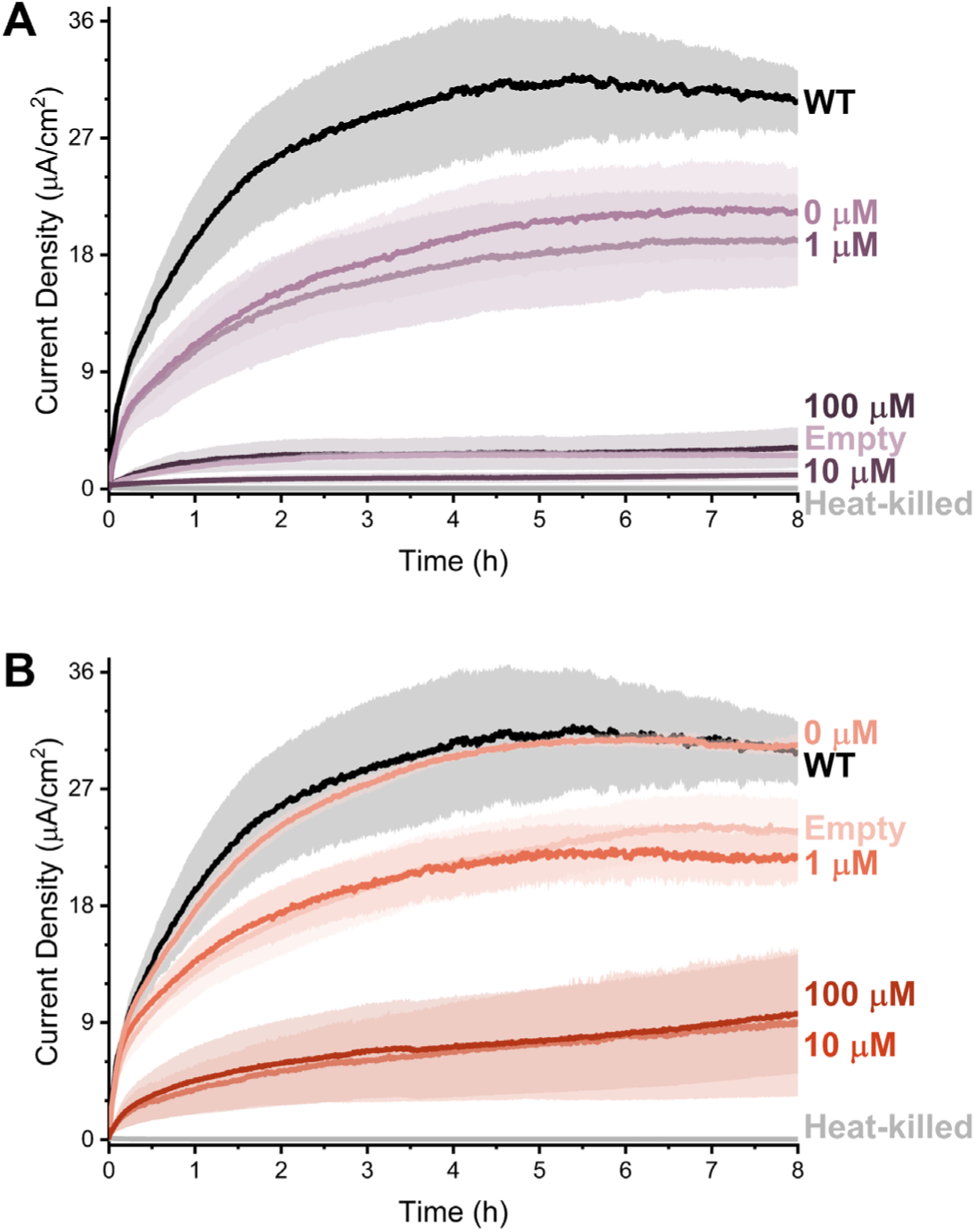
Overexpression of either *mtrA* or *mtrC* impairs electrode reduction. **(A)** Chronoamperometry of live and heat-killed WT cells compared to Δ*mtrA* + LP2::*cymR* + LP1::P_cymRD_-*mtrA* cells grown in different concentrations of cumate (0 µM, 1 µM, 10 µM, or 100 µM), as well as to Δ*mtrA* + LP2::*cymR* + LP1::P_cymRD_-empty cells (0 µM cumate), shows that the induction level of *mtrA* alters current production. **(B)** Chronoamperometry of live and heat-killed WT cells compared to Δ*mtrC* + LP2::*cymR* + LP1::P_cymRD_-*mtrC* cells grown in different concentrations of cumate (0 µM, 1 µM, 10 µM, or 100 µM), as well as to Δ*mtrC* + LP2::*cymR* + LP1::P_cymRD_-empty cells (0 µM cumate), shows that the induction level of *mtrC* alters current production. Each curve represents the mean of three independent biological replicates and the error bars indicate standard error of the mean. The BESs in which chronoamperometry measurements were made did not contain any cumate.

The same experimental design was used to evaluate the relationship between MtrC expression level and electrode reduction. The current density produced by the Δ*mtrC*-Empty Cassette strain was significantly lower than WT but had a notably greater magnitude than the other single gene deletion strains (max. 23 μA/cm²) **(Figure 4B)**. The muted defect of *mtrC* deletion was eliminated by complementation of *mtrC* with 0 µM cumate, which restored electrode reduction to a magnitude nearly identical to WT. As with MtrA, the level of MtrC expression enabled by basal expression from CymR-repressed P_cymRD_ was sufficient to boost current above the level created by deletion of the respective decaheme cytochrome. The current density produced by 1 µM cumate induction of MtrC expression was similar to that produced by Δ*mtrC*-Empty Cassette. Similarly to *mtrA*, induction of *mtrC* with 10 µM or 100 µM cumate resulted in current densities much lower than those produced by WT. Yet, heme visualization of these highly induced cells showed a prominent band at the expected MW for MtrC. **(Supplemental Figure 6A-C)**. Deletion and complementation of MtrC reveals that, although MtrC contributes to electrode reduction, it is not strictly required for *V. natriegens* to transfer electrons to the electrode. The recovery of the WT electrode reduction phenotype by complementation demonstrates that the minor electrode reduction defect of Δ*mtrC V. natriegens* is caused by the absence of the *mtrC* gene product, not by disruption of the remaining *mtrCAB* operon sequence. Notably, overexpression of either decaheme cytochrome produces a profoundly deleterious effect on electrode reduction, despite visualization of the overexpressed decaheme in the cell lysates.

### High induction of *mtrA* or *mtrC* expression disrupts the balance of cytochromes required for EET

To better interrogate why overexpression of either of the decaheme cytochromes hinders electrode reduction, we sought to assess how this overexpression impacts the expression of CymA, PdsA, and the other decaheme cytochrome. To do so, we first sought to identify a band in the ECL analysis of whole cell lysates that could be attributed to CymA (predicted MW: 25.050 kDa) or PdsA (predicted MW: 20.804 kDa). Comparison of WT to the double knock-out Δ*cymApdsA* revealed the loss of a subtle band in the region of 20-25 kDa **(Supplemental Figure 7A)**. However, this band was very difficult to discern because it was sandwiched between two prominent bands **(Supplemental Figure 7B)**. To improve our ability to visualize the CymA/PdsA dependent band, we deleted PN96_RS12985 (predicted MW: 21.113 kDa, 82.93% identity to *Vibrio cholerae* Cytochrome c_4_ (47)). Deleting PN96_RS12985 resulted in the loss of a ∼22 kDa band, which allowed clear visualization of the CymA/PdsA band **(Supplemental Figure 7A)**. Comparison of triple knock-out Δ*cymApdsA*PN96_RS12985 cells to the single ΔPN96_RS12985 deletion enabled clear recognition of heme signal attributable to a combination of CymA and PdsA **(Supplemental Figure 7B)**.

The PN96_RS12985 deletion enables visualization of a single band that is lost when both *cymA* and *pdsA* are deleted. We investigated whether this band is the product of CymA, PdsA, or a combination of both proteins by making single deletions of *cymA* and *pdsA* within a ΔPN96_RS12985 background and examining the effect of those deletions on the cytochrome *c* pattern produced by ECL analysis. We observed that both single deletion strains produced a band that was absent in the double deletion **(Supplemental Figure 7C)**. Based on this finding, we conclude that this band represents both cytochromes, CymA and PdsA.

By using the Cytochrome *c*_4_ (PN96_RS12985) deletion background to better visualize the combined expression of CymA and PdsA, we aimed to test the effect of overexpressing MtrA or MtrC on the expression level of CymA/PdsA and the other decaheme cytochrome. For both the *mtrA*-expressing strain and the *mtrC*-expressing strain, we compared biological replicates of cells cultured with 1 µM cumate to WT cells and to cells cultured with a higher concentration of cumate. At low levels of induction (1 µM cumate) of *mtrA* expression, a band corresponding to MtrA, a band corresponding to CymA+PdsA, and a band corresponding to MtrC were all visible **(Figure 5A)**. This pattern of cytochromes *c* closely resembled that of WT cells. However, at higher levels of *mtrA* induction (10 µM cumate), the intensity of the bands corresponding to CymA+PdsA and MtrC was greatly diminished while a band attributable to MtrA was nonetheless prominent **(Figure 5A, Supplemental Figure 8A-C)**. A similar trend was observed for varied induction of *mtrC*. At low levels of induction (1 µM cumate), bands corresponding to MtrC, CymA+PdsA, and MtrA were all detected, as was observed from WT cells **(Figure 5B)**. Higher induction of *mtrC* (100 µM cumate) greatly diminished the intensity of the bands corresponding to CymA+PdsA and MtrA, despite the presence of a prominent MtrC band **(Figure 5B, Supplemental Figure 8D-F)**. Taken together, these observations indicate that overexpression of either *mtrA* or *mtrC* impairs expression of the other cytochromes involved in EET and that a comparably narrow range of expression levels is needed to accumulate *mtrA* or *mtrC* without impairing the production of other EET cytochromes *c*. These findings strongly suggest that the negative effect on electrode reduction of high levels of cumate induction of decaheme cytochrome expression is caused by disruption of the balance of multiheme cytochromes required for EET.

**Figure 5:**
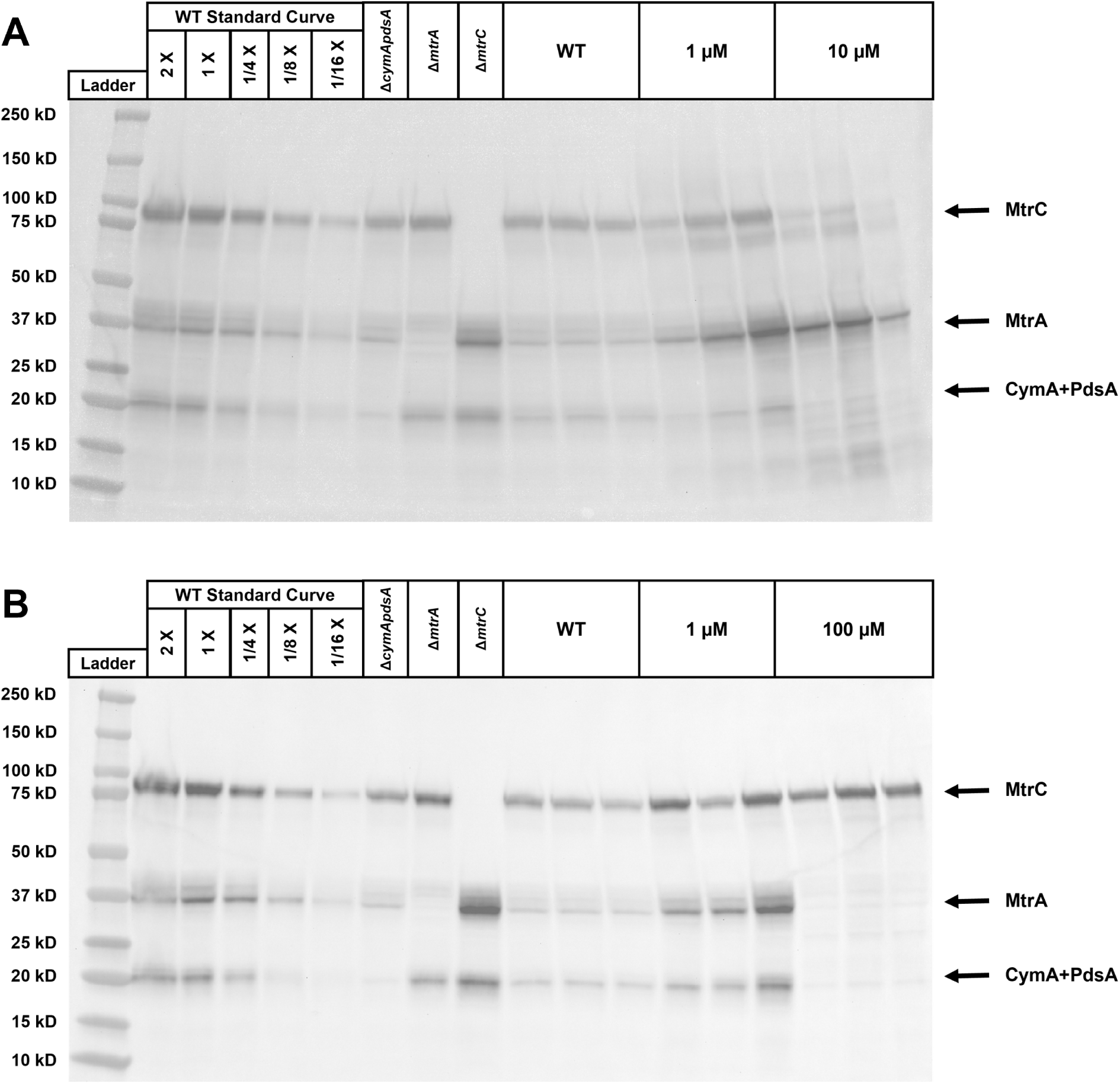
Overexpression of either *mtrA* or *mtrC* impairs expression of other cytochromes. **(A)** Comparison of the band patterns produced by ECL of whole cell lysates of WT cells and cells containing an inducible copy of *mtrA*, induced with 1 µM or 10 µM cumate (three biological replicates each), reveals loss of band intensity for MtrC and CymA/PdsA under the 10 µM cumate condition. **(B)** Comparison of the band patterns produced by ECL of whole cell lysates of WT cells and cells containing an inducible copy of *mtrC*, induced with 1 µM or 100 µM cumate (three biological replicates each), reveals loss of band intensity for MtrC and CymA/PdsA under the 100 µM cumate condition. A standard curve of WT cells loaded at five different concentrations demonstrates that the observed band intensities are within a discernible range. Wells showing the band pattern produced by Δ*cymApdsA*, Δ*mtrA*, and Δ*mtrC* allow attribution of individual bands to CymA+PdsA, MtrA, and MtrC, respectively. All strains carry a PN96_RS12895 deletion in addition to the other stated genotypic features.

## Discussion

Here, we investigated the role of the outer membrane protein *mtrB* and the multiheme cytochromes *cymA*, *pdsA*, *mtrA*, and *mtrC* in electrode reduction by *V. natriegens*. We find that each of these genes contributes to electrode reduction. While single gene deletion of each of the other genes largely eliminates electrode reduction, deletion of *mtrC* produces only a modest loss of electron transfer to the electrode. We observed that differences in EET activity did not result in significant growth effects. Most intriguingly, we uncovered that for *V. natriegens*, carefully balanced cytochrome *c* expression is required for electrode reduction.

Our results expand our understanding of how aspects of electrode reduction in *V. natriegens* share similarities with iron reduction in *V. natriegens* and electrode reduction in *S. oneidensis*. Both gluconate and glucose support electrode reduction by *V. natriegens*, a finding that differs from a published study on iron reduction, which found robust iron reduction with gluconate but minimal iron reduction with glucose (18). With gluconate as the electron donor, we routinely observe a maximum current density of around 25-40 µA/cm^2^ from *V. natriegens* under our experimental conditions. When studies observe different organisms under different electrolyte and electron donor conditions, equivalent comparisons of current magnitudes are difficult. However, we recently measured a maximum current density of around 6.2 µA/cm^2^ from *S. oneidensis* using similar BES protocols to those used here (48), suggesting that the magnitude of *V. natriegens* electrode reduction is comparable with that of *S. oneidensis*. The *S. oneidensis* Mtr pathway supports both iron reduction and electrode reduction (12). Our data indicate that this is true for *V. natriegens* as well, because our findings on the genetic requirements for electrode reduction agree with the findings of Conley, *et al*. regarding iron reduction (18).

Surprisingly, we observe substantial electrode reduction in the absence of MtrC, raising the question of how electrons reach the electrode in the absence of MtrC. In *S. oneidensis*, the EET phenotype of MtrC can be supported by other multiheme cytochromes localized to the cell surface when *mtrC* alone is deleted (49). However, flux through an alternative cell surface electron transfer protein seems unlikely to occur in *V. natriegens*, where *mtrC* appears to be the only homolog of *S. oneidensis* extracellular multiheme cytochromes. Notably, MtrAB of *S. oneidensis* can be modified to support iron reduction in the absence of MtrC by making only two point mutations (50). Perhaps the MtrAB of *V. natriegens* has evolved a greater ability to reduce solid electron acceptors than the WT MtrAB of *S. oneidensis*. Alternatively, MtrC-independent electrode reduction might be explained by electron shuttling. MtrA might reduce a soluble electron acceptor that is, in turn, oxidized by the electrode. Such a mediator may be biotic in origin; *S. oneidensis* can utilize endogenously produced flavin and quinone molecules as electron shuttles (51, 52). *V. natriegens* possesses genes predicted to enable the biosynthesis of these molecules, but whether *V. natriegens* releases these molecules and can utilize them as mediators remains to be investigated. Another possibility is that MtrC-independent electrode reduction might be facilitated by metal ions present in the medium used in the BES. Of particular note is the 51 µM ferric chloride. Some studies of *S. oneidensis* iron reduction have reported reduction of soluble iron by cells lacking outer membrane cytochromes, and soluble iron has previously been suggested to facilitate electron transfer between an electrode and *S. oneidensis* cells (32, 53, 54). Further investigation will be needed to unravel how *V. natriegens* performs robust electrode reduction without the need for MtrC. Elucidating this mechanism may also help explain why non-essential *mtrC* has been retained in the *V. natriegens* genome, an especially interesting question in light of the deleterious consequences of its overexpression.

We observed that without tight control, expression of either MtrA or MtrC diminishes production of other multiheme cytochromes and results in an EET-impaired phenotype. Interestingly, although we see strong deleterious effects of decaheme cytochrome overexpression, we do not observe similar electrode reduction defects from the presumably strong unrepressed P_cymRC_ expression of CymA, PdsA, and MtrB. However, without quantification of transcription and translation levels of each cytochrome, we cannot be certain how the production rates of CymA, PdsA, and MtrB compare to those of MtrA and MtrC. Nonetheless, the absence of an observation of a ceiling on induction for *cymA*, *pdsA*, and *mtrB* suggests some possible explanations for the strict limits on induction of *mtrA* and *mtrC*. We speculate as to a few potential reasons the balance of multiheme cytochromes may be so exquisitely sensitive to decaheme cytochrome overexpression.

For one, the decaheme cytochromes may impose a greater burden on the supply of biosynthesized heme and the capacity of the cytochrome *c* maturation system than the tetraheme CymA or the diheme PdsA. Limits on multiheme cytochrome *c* production have been found to constrain expression of EET pathways ported into heterologous chassis (37, 38). We may be encountering similar limitations when overexpressing multiheme cytochromes *c* within *V. natriegens*. Other work in *S. oneidensis* has observed that eliminating some periplasmic cytochromes can enhance EET (55, 56). If the total *c*-type cytochrome production capacity of a bacterium is constrained, we would expect to observe the types of tradeoffs between expression of different cytochromes that were observed in *S. oneidensis* and in our data. Alternatively, the ceiling on productive decaheme cytochrome expression might arise from the constraints of forming the porin:cytochrome complex. Interdependency of proteins within the MtrCAB complex has been described in *S. oneidensis* — MtrA plays a critical role in protecting MtrB from degradation (43). Thus, preserving proper stoichiometry of the MtrCAB complex components may be essential for stable maintenance of EET phenotype. Our highly induced decaheme cytochrome expression may dysregulate the stoichiometry of the complex, triggering as-of-yet undefined stress responses or regulatory mechanisms. However, any such explanation must contend with the apparent lack of EET defect when MtrB expression was unconstrained.

By characterizing the cytochrome balance required for current production by *V. natriegens*, this work identifies design parameters and constraints for the development of *V. natriegens* for bioelectronic applications. Our findings suggest that the expression level of *cymA*, *pdsA*, *mtrB*, or *mtrA* could be used to exercise transcriptional control of the electronic output of *V. natriegens*. By contrast, our results indicate that modulation of *mtrC* expression is likely to yield significantly lower dynamic range compared to modulation of the other four genes. Additionally, we find evidence suggesting that high levels of expression of *cymA* and *pdsA* do not impair *V. natriegens* EET or anaerobic growth, making these two genes more appealing targets for synthetic biological manipulation than the more tightly constrained expression of *mtrA*. Some biotechnological applications may benefit from maximizing the EET flux in *V. natriegens*. However, none of our strains overexpressing single EET genes produced greater current than WT, suggesting that none of CymA, PdsA, or MtrCAB individually constitute a rate-limiting bottleneck. Perhaps increasing the expression level of multiple EET pathway components could increase current production above WT levels. However, our observations indicate that expression of MtrA or MtrC much beyond WT levels is deleterious for EET. Identification of the mechanism underlying this ceiling on productive decaheme cytochrome expression might illuminate paths to enhancing *V. natriegens* EET activity. In all, our characterization of the cytochrome balance required to support electrical current production by *V. natriegens* provides key insights for the future design of engineered microbial bioelectronic technologies using *V. natriegens* as a novel chassis.

## Materials and Methods

### Media

Cultivation of *E. coli* was performed in liquid LB, which was prepared from “LB Broth (Miller)” from Sigma (L3522), and on LB-agar plates, which were prepared from “LB Broth with agar (Miller)” from Sigma (L3147). Multiple rich media formulations were used for cultivation of *V. natriegens*. LBv2 was prepared from “LB Broth (Miller)” from Sigma (L3522) supplemented with 204 mM additional NaCl, 4.2 mM KCl, and 23.14 mM MgCl_2_. LBv2-agar plates were prepared from “LB Broth with agar (Miller)” from Sigma (L3147) supplemented with 204 mM additional NaCl, 4.2 mM KCl, and 23.14 mM MgCl_2_. BHIv2 was prepared from “Brain Heart Infusion Broth” from Sigma (53286) supplemented with 204 mM additional NaCl, 4.2 mM KCl, and 23.14 mM MgCl_2_. ZYM-5052-v2 (36) was composed of 10 g/L N-Z-amine A (Millipore C0626), 5 g/L yeast extract (Millipore 70161), 25 mM Na_2_HPO_4_, 25 mM KH_2_PO_4_, 50 mM NH_4_Cl, 5 mM Na_2_SO_4_, 2 mM MgSO_4_, 5 g/L glycerol, 0.5 g/L glucose, 2 g/L α-lactose monohydrate, 204 mM NaCl, 4.2 mM KCl, 23.14 mM MgCl_2_, and trace metals. The final concentrations of trace metals in ZYM-5052-v2 were 10 µM FeCl_3_, 4 µM CaCl_2_, 2 µM MnCl_2_, 2 µM ZnSO_4_, 400 nM CoCl_2_, 400 nM CuCl_2_, 400 nM NiCl_2_, 400 nM Na_2_MoO_4_, 400 nM Na_2_SeO_3_, and 400 nM H_3_BO_3_. Trace metals were prepared as a combined stock solution at 500 times the aforementioned concentrations in 6 mM HCl. BES measurements were performed in *Vibrio natriegens* Extracellular Electron Transfer Medium (VEETM), which was composed of 93.05 mM Na_2_HPO_4_, 48.4 mM KH_2_PO_4_, 439.9 mM NaCl, 22.1 mM NH_4_Cl, 6.61 mM MgSO_4_, 570 µM CaCl_2_, 171 µM EDTA, 51 µM FeCl_3_, 6.2 µM ZnCl_2_, 1.6 µM H_3_BO_3_, 970 nM CuCl_2_, 420 nM CoCl_2_, and 99 nM MnCl_2_. Chloramphenicol (Sigma C0378) was added to all media used to grow cells carrying plasmids derived from pST140 (44) (except where otherwise noted). A final chloramphenicol concentration of 34 µg/mL was used when culturing *E. coli* with both solid and liquid media. For *V. natriegens* cultures, 2 µg/mL chloramphenicol was used in solid media, while 4 µg/mL was used in liquid media.

### Culturing Methods

To prepare cells for experiments, a small amount of frozen glycerol stock was streaked onto LBv2-agar plates and incubated for 12 hours at 37 °C. From these plates, single colonies were used to inoculate 3 mL liquid cultures of LBv2 medium in 15 mL culture tubes, which were incubated for 12 hours at 37 °C with 250 rpm shaking to yield dense, stationary phase cultures.

For BES experiments, which require large quantities of cells, 50 µL of the dense, stationary phase LBv2 culture was added to 50 mL of ZYM-5052-v2 in a 250 mL flask. Following the medium, incubation temperatures, and incubation durations of Fuchs, *et al.*, (36) which produced robust *c*-type cytochrome expression in *V. natriegens*, flask cultures were shaken at 250 rpm at 30 °C until the cell density reached OD_600_ = 0.6 (corresponding to log phase growth, approximately 4.5 hours after inoculation). When the culture reached an OD_600_ = 0.6, the incubation temperature was decreased to 25 °C and cumate (4-isopropylbenzoic acid, Sigma, 268402) was added at indicated concentrations when specified to induce *mtrA* or *mtrC* expression. Cells were harvested after 24 hours of total incubation in ZYM-5052-v2.

For SDS-PAGE + Enhanced Chemiluminescence experiments, which require fewer cells, smaller-scale cultures were grown by adding 2 µL of LBv2 culture to 2 mL of ZYM-5052-v2 in a 24-well deep-well plate and incubating at 550 rpm (Labnet Vortemp 56, 3 mm orbit) and 30 °C until the cell density reached OD_600_ = 0.6, approximately 4.5 hours later. When the culture reached OD_600_ = 0.6, the incubation temperature was decreased to 25 °C and cumate was added at indicated concentrations when specified to induce *mtrA* or *mtrC* expression. Cells were harvested after 24 hours of total incubation in ZYM-5052-v2.

### Strains and Plasmids

All strains were derived from *Vibrio natriegens* ATCC 14048. All strains were stored at -80 °C as glycerol stocks. Glycerol stocks were prepared by combining 700 µL of 50% (v/v) glycerol with 700 µL of an LBv2 culture grown for 6-8 h at 37 °C with 250 rpm shaking, following the method described in Stukenberg, *et al.* (44). **Supplemental Table 1** lists all strains of *V. natriegens* used in this chapter and the process used to create each strain.

Genomic deletions and integrations of new gene expression cassettes were performed using the NT-CRISPR method described by Stukenberg, *et al.* (44). This approach combines the natural transformation of *V. natriegens* by linear dsDNA molecules (herein referred to as tDNA) with CRISPR-Cas9 counterselection. NT-CRISPR is facilitated by plasmid-encoded expression of isopropyl β-D-1-thiogalactopyranoside (IPTG)-inducible *tfoX* expression (required to induce natural competency for tDNA uptake) and anhydrotetracycline (aTc)-inducible Cas9 and gRNA expression (for counter-selection).

### Selection of sites for chromosomal integration

Landing Pad 1 (LP1) was used for chromosomal integration of gene expression cassettes encoding the complementary expression of *V. natriegens* EET genes. To identify a suitable locus for chromosomal integration, we used the permissR computational tool to identify large intergenic regions that did not overlap possible insertion elements (57). Informed by the analysis from permissR, we selected the 1,500 bp intergenic region between genes PN96_RS17100 and PN96_RS17105 on chromosome 2 as LP1. The new genetic material was integrated between the nucleotides located 765 and 766 nt upstream of the 5’ end of the CDS for PN96_RS17100 (the site is between 735 and 736 nt downstream of the 3’ end of the CDS for PN96_RS17105).

Landing Pad 2 (LP2) was used for chromosomal integration of a gene expression cassette encoding the constitutive expression of the transcription factor CymR. This locus was selected as an integration site based on its use by Stukenberg, *et al.* (44) LP2 lies within a 729 bp intergenic region between the genes PN96_RS06225 and PN96_RS06230 on chromosome 1. The new genetic material was integrated between the nucleotides located 491 and 492 nt downstream of the 3’ end of the CDS for PN96_RS06225 (the site is between 238 and 239 nt downstream of the 3’ end of the CDS for PN96_RS06230).

### Assembly of NT-CRISPR plasmids

In NT-CRISPR, the CRISPR-Cas9 counter-selection requires expression of a gRNA targeting a genomic DNA sequence that will be disrupted by the desired deletion or insertion. For each unique genomic locus targeted for deletion or insertion, we assembled a unique NT-CRISPR plasmid carrying a gRNA specific to the target locus. These gRNA sequences were designed using the Benchling sgRNA design tool. Plasmids expressing the new gRNA sequences were assembled from pST140 (44) and a short gRNA-encoding dsDNA fragment produced by annealing two single-stranded oligonucleotides. gRNA and oligonucleotide sequences for each modified locus are listed in **Supplemental Table 2**. For each plasmid, the two oligonucleotides were annealed by mixing 1.25 μL of each oligonucleotide (100 μM) in 7.5 μL of 10 mM Tris buffer (Sigma, T6066) and heating with a descending temperature gradient from 94 °C to 37 °C, with the temperature decreased 0.1 °C every five seconds, thereby forming a dsDNA molecule consisting of the gRNA sequence with additional 4-bp 5’ and 3’ overhangs designed for assembly into pST140 by Golden Gate assembly. To prepare these dsDNA fragments for ligation into the pST140-derived backbone, the dsDNA fragments were phosphorylated by preparing 50 µL reactions consisting of 10 µL annealing sample, 5 µL of 10X T4 Ligase Buffer (New England Biolabs), 0.625 µL of T4 PNK enzyme (New England Biolabs), and 34.375 µL of water. Phosphorylation reactions were incubated at 37 °C for 30 minutes.

New plasmids were assembled through Golden Gate reactions between the linear inserts described above and pST140, performed in T4 Ligase Buffer (New England Biolabs) using BsaI-HF (New England Biolabs) and T4 Ligase (New England Biolabs). After completion of the Golden Gate reactions, the reaction mixture was transformed into competent NEB 10β *E. coli* cells, which were plated on LB-agar with 34 µg/mL chloramphenicol to isolate clonal plasmid populations. Colonies carrying a successful assembly product were identified by selecting colonies without sfGFP expression, because the sfGFP expression cassette on pST140 should be removed during a successful BsaI-mediated Golden Gate assembly. Samples of purified clonal plasmid populations were obtained by culturing individual colonies for 12-16 hours in LB at 3 °C with 250 rpm shaking and extracting plasmid DNA using a Monarch Plasmid Miniprep Kit (New England Biolabs). All plasmids were fully sequenced through Oxford Nanopore sequencing by Plasmidsaurus. Full plasmid sequences are listed in **Supplemental Table 3**.

### Transformation of NT-CRISPR plasmids into V. natriegens

Before making a chromosomal modification in a given strain, that strain must be transformed with the NT-CRISPR plasmid carrying a proper gRNA sequence for the desired chromosomal modification. Electroporation was utilized to introduce NT-CRISPR plasmids into *V. natriegens* strains. To prepare electrocompetent cells, a small amount of frozen glycerol stock was streaked onto LBv2-agar plates and incubated for 12 hours at 37 °C. From these plates, single colonies were used to inoculate 3 mL of liquid LBv2 medium in 15 mL culture tubes, which were incubated for 12 hours at 37 °C with shaking at 250 rpm. Fifty microliters of these LBv2 precultures were used to inoculate 50 mL BHIv2 cultures in 250 mL flasks. Flask cultures were incubated at 37 °C with 250 rpm shaking until the OD_600_ reached 0.5, at which point cells were cooled on ice for 15 minutes, then harvested by centrifugation at 4000 x g for 20 minutes at 4 °C. After removing the supernatant, the cell pellets were washed three times by resuspending the cells in 40 mL of ice-cold sterile Electroporation Buffer (680 mM sucrose, 7 mM K_2_HPO_4_, pH 7), pelleting the cells by centrifugation at 4000 x g for 15 minutes at 4 °C, and disposing of the supernatant. Finally, the cells were resuspended to OD_600_ = 5-10 with Electroporation Buffer. These cell suspensions were stored in 50 µL aliquots at -80 °C until electroporation.

For transformation by electroporation, 100 ng of plasmid DNA was mixed with a 50 µL aliquot of electrocompetent *V. natriegens* cells in a 1 mm gap electroporation cuvette (Bio-Rad, 1652089). Electroporation was performed at 0.9 kV using a MicroPulser Electroporator (Bio-Rad). Cells were recovered for one hour at 37 °C, shaking at 250 rpm, in Recovery Medium (BHIv2 with 680 mM sucrose) prior to plating onto LBv2-agar plates supplemented with 2 µg/mL chloramphenicol. After 12 hours of incubation at 37 °C, individual colonies were used to inoculate 3 mL LBv2 liquid cultures to be used to prepare glycerol stocks.

### Natural transformation with linear tDNA

To produce ample DNA for natural transformation, linear tDNA molecules (**Supplemental Table 4**) were amplified by polymerase chain reaction (PCR) using Q5 polymerase (New England Biolabs) and column purified using “Econospin Spin Columns for DNA” (Epoch) and DNA cleanup buffers from Qiagen (Buffer PB, Buffer PE). All tDNAs included 3kb of homology to the sequence of the chromosome on both sides of either the sequence to be deleted or the locus for insertion. For tDNAs facilitating insertion of new sequences into the chromosome, the 3 kb homology arms were placed on either side of the sequence of genetic parts **(Supplemental Table 5)** to be inserted.

To prepare cells for natural transformation, a small amount of frozen glycerol stock of NT-CRISPR plasmid-containing cells was streaked onto LBv2-agar plates and incubated for 12 hours at 37 °C. From these plates, single colonies were precultured for 16-17 h at 30 °C with 250 rpm shaking in 15 mL culture tubes containing 3 mL of LBv2 supplemented with 4 µg/mL chloramphenicol and 100 µM IPTG (added to induce *tfoX* expression and thereby trigger natural competence in *V. natriegens*). Natural transformation was initiated by mixing 3.5 µL of preculture and 10 ng of tDNA with 350 µL of Sea Salt Medium (28 g/L “Sea salts”, S9883, Sigma) supplemented with 100 µM IPTG and incubating statically at 30 °C for 5 h. After the incubation, expression of Cas9 and the gRNA were induced by adding 1 mL of antibiotic-free LBv2 supplemented with 200 ng/mL aTc and incubating at 30 °C with 250 rpm shaking for 1 h. Cells were subsequently plated onto LBv2-agar plates supplemented with 2 µg/mL chloramphenicol and 200 ng/mL aTc to select for cells in which successful natural transformation by the tDNA resulted in the elimination of the native chromosomal sequence targeted by the plasmid-expressed gRNA. After 12 hours of growth on the plates, individual colonies were screened to confirm that the desired chromosomal modification was successful. The genomic locus of interest was PCR amplified using primers binding approximately 100 bp upstream and downstream of the outer limits of the tDNA **(Supplemental Table 6)**. PCR amplicons were fully sequenced through Oxford Nanopore sequencing by Plasmidsaurus to validate the chromosomal modifications contained within each strain **(Supplemental Table 1)**. Sequence-validated strains were stored at -80 °C in glycerol stocks.

### Plasmid curing

To remove the NT-CRISPR plasmid from strains with completed chromosomal modifications, cells were grown in 3 mL of LBv2 without antibiotics in 15-mL culture tubes at 37 °C with 250 rpm shaking for 6–7 hours. A sterile inoculating loop was used to streak samples of these dense liquid cultures onto antibiotic-free LBv2 agar plates to isolate single colonies. After overnight incubation at 37 °C, colonies were screened for loss of the plasmid by re-plating on both antibiotic-free LBv2-agar and LBv2-agar supplemented with 2 µg/mL chloramphenicol. Colonies that grew only on antibiotic-free plates were considered plasmid-cured. Verified plasmid-cured strains were preserved as glycerol stocks at -80 °C.

### Bioelectrochemical System measurements

Measurements of electrode reduction were performed in three electrode BESs with two chambers separated by a cation exchange membrane (CMI-7000, Membranes International). The cathodic chamber contained a 1 mm diameter titanium wire cathode and 115 mL of VEETM. The anodic chamber contained a Ag/AgCl (3 M KCl) reference electrode (CHI111, CH Instruments) and a working anode consisting of a cylinder of carbon felt (Alfa Aesar) with a geometric surface area of 22 cm^2^ suspended from a 1 mm diameter titanium wire. Prior to beginning electrochemical measurements, the anodic chamber was filled with 111 mL of VEETM, continuously mixed with a stir bar at 320 rpm, maintained at 30 °C via water jacketing, and continuously sparged with N_2_ gas to maintain an anaerobic environment. To measure electrode reduction, the working electrode was polarized at +200 mV vs. the Ag/AgCl reference electrode and the current, averaged over every 36 seconds, was recorded using a VMP-300 or VSP-300 potentiostat (Bio-Logic). Measurements of cell-free VEETM were recorded prior to the addition of cells.

To prepare cells for injection into the BES, cells were pelleted, washed to remove any residual media, and resuspended at a high cell density. Specifically, cells from 50 mL ZYM-5052-v2 cultures were first pelleted by centrifugation at 4000 x g and 4 °C for 15 minutes and the supernatant was discarded. The retained cells were washed two times, where each wash step consisted of resuspending the cell pellets in VEETM (using 40% of the original volume), pelleting the cells via centrifugation at 4000 x g and 4 °C for 15 minutes, and decanting the supernatant. After the second wash, each cell pellet was resuspended in VEETM to yield an OD_600_ = 20. To generate heat-killed controls, these dense cell suspensions were heated to 70 °C for one hour. Three milliliters of washed cell suspensions and 6 mL of 20X electron donor solution were simultaneously injected into BESs, yielding an initial cell density of OD_600_ = 0.5. The 20X electron donor solution was 1.1 M sodium gluconate in VEETM (final concentration = 55 mM) for all BES experiments, except experiments comparing different electron donors, in which 1.1 M glucose in VEETM (final concentration = 55 mM) and 1X VEETM were also used.

### Growth curve measurement and analysis

To monitor growth dynamics during BES operation, the medium in the anodic channel was periodically sampled at approximately three-hour intervals for 24 hours after the addition of cells to the BES. At each sampling time, 1 mL of medium was removed from each BES. This single sample was split into technical triplicates for five-fold dilution into VEETM and subsequent OD_600_ measurement. The mean OD_600_ values were used to fit a microbial growth curve for each BES. To model microbial growth curves, we used the Zwietering-modified Gompertz equation **(1)** (58).

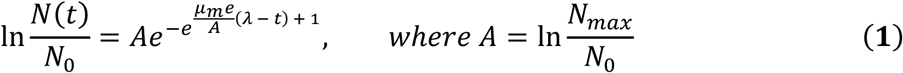

In this equation, t is time, N(t) is the cell number at time t, N_0_ is the initial cell number, N_max_ is the cell number in stationary phase, μ_m_ is the maximal growth rate, and λ is the duration of lag phase (58). We rearranged Equation 1 to describe N(t) in terms of N_0_, N_max_, μ_m_, and λ **(2)**.

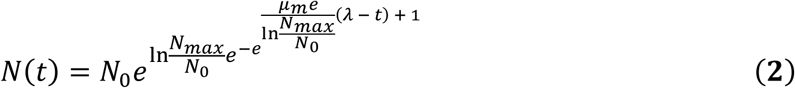

We assumed that OD_600_ was linearly proportional to cell number over the range of our OD_600_ measurements and fit Equation 2 for N(t) to each time series of OD_600_ measurements using the Matlab function “fitnlm” and the initial parameters: N_0_ = 0.5, N_max_ = 1.1, μ_m_ = 0.07, and λ = 6. To test whether any of the strains showed significant differences from any other strain in any of the three parameters N_max_, μ_m_, and λ, we performed a random block ANOVA, based on our randomized block experimental design (59). ANOVA analysis was performed using the R script from Zweifach (60).

### SDS-PAGE and Enhanced Chemiluminescence

Evaluation of the cytochrome *c* content of *V. natriegens* cells was performed through SDS-PAGE of whole cell lysates, transfer to a nitrocellulose membrane, and treatment of the membrane with Clarity Western Enhanced Chemiluminescence Substrate (Bio-Rad). The luminol-based Enhanced Chemiluminescence (ECL) reagent produces light in response to the peroxidase activity of the covalently bound heme cofactors of *c*-type cytochromes. To prepare cells for heme blots, 2 mL ZYM-5052-v2 cultures were centrifuged for 20 minutes at 4000 x g and 4 °C, and the cell pellets were resuspended to a cell density of OD_600_ = 30 in 1X PBS. For Supplemental Figures 5 & 6, cell suspensions at OD_600_ = 20 in VEETM were used instead, so as to evaluate the cytochromes *c* of the cell suspensions that were used to inoculate BESs in Figure 4A & 4B. In all cases, the cell suspensions were combined 1:1 with a solution consisting of 20 parts 2X Laemmli Sample Buffer (Bio-Rad, 1610737) and 1 part β-mercaptoethanol. The resulting mixture was incubated at 96 °C for 15 minutes to lyse the cells.

Thirty microliters of cell lysate was loaded into each well of a 4-20% Criterion TGX Stain-Free Protein Gel (Bio-Rad, 5678094) that was run at 300 V in 1X Tris/Glycine/SDS running buffer (Bio-Rad, 1610732) until the lowest mass band of the Precision Plus Protein Dual Color Standards ladder (Bio-Rad, 1610374) reached the bottom of the gel. The separated protein bands were transferred from the gel to a nitrocellulose membrane (Bio-Rad, 1704159) using a Trans-Blot Turbo Transfer System (Bio-Rad). The nitrocellulose membrane was treated with 7 mL of ECL Substrate (3.5 mL of “luminol/enhancer solution” + 3.5 mL of “peroxide solution”) for four minutes, after which the luminescence was measured with a six-minute exposure on a ChemiDoc Go Imaging System (Bio-Rad).

Image analysis was performed to extract band volumes to confirm the qualitative conclusions drawn from the blot images. First, rolling ball background subtraction was performed in ImageJ using a radius of 75 pixels. Next, lane boundaries were manually identified. The upper and lower boundaries of the bands corresponding to CymA+PdsA, MtrA, and MtrC were determined automatically by comparing the lanes from WT samples to those from Δ*cymApdsA*, Δ*mtrA*, and Δ*mtrC*, respectively. Specifically, traces plotting the average pixel intensity as a function of migration distance for the WT lanes were subtracted from the corresponding traces for the deletion strain lanes; the largest resulting peak was assigned to the deleted cytochrome. These computationally defined upper and lower band boundaries were then combined with the manually defined lane boundaries to define a box containing each band of interest in each well. Band volumes were computed by summing the intensities of all of the pixels within these boxes.

## Acknowledgements

We thank Xinyue Luo for technical assistance measuring the role of MtrB in electrode reduction. We thank Swetha Sridhar for assistance with the computational identification of a suitable locus for chromosomal insertion in *V. natriegens* (site LP1). We thank Shyam Bhakta for sharing the P_cymRD_ promoter part. M.D.C. acknowledges the Cancer Prevention and Research Institute of Texas (award # RR1900063) for primary financial support, as well as the U.S. Army Research Office (Multidisciplinary University Research Initiative, award # W911NF2210239), and the National Science Foundation (SemiSynBio, award # 2227526; NSF Research Traineeship, award # 1828869) for supplemental financial support. W.-C. C. acknowledges the financial support of the National Science Foundation (Emerging Frontiers in Research and Innovation, award # 2223678). C. M. A.-F. acknowledges financial support by the National Science Foundation (Emerging Frontiers in Research and Innovation, award # 2223678; SemiSynBio, award # 2227526) and the U.S. Army Research Office (Multidisciplinary University Research Initiative, award # W911NF2210239).

## Notes

### Competing Interest Statement

The authors have declared no competing interest.

## References

1. Gralnick JA, Newman DK. 2007. Extracellular respiration. Molecular Microbiology 65:1–11.

2. Atkinson JT, Chavez MS, Niman CM, El-Naggar MY. 2023. Living electronics: A catalogue of engineered living electronic components. Microbial Biotechnology 16:507–533.

3. Shi L, Dong H, Reguera G, Beyenal H, Lu A, Liu J, Yu H-Q, Fredrickson JK. 2016. Extracellular electron transfer mechanisms between microorganisms and minerals. Nat Rev Microbiol 14:651–662.

4. Myers CR, Nealson KH. 1988. Bacterial Manganese Reduction and Growth with Manganese Oxide as the Sole Electron Acceptor. Science 240:1319–1321.

5. Hunt KA, Flynn JM, Naranjo B, Shikhare ID, Gralnick JA. 2010. Substrate-Level Phosphorylation Is the Primary Source of Energy Conservation during Anaerobic Respiration of Shewanella oneidensis Strain MR-1. Journal of Bacteriology 192:3345–3351.

6. Lovley DR, Phillips EJP. 1988. Novel Mode of Microbial Energy Metabolism: Organic Carbon Oxidation Coupled to Dissimilatory Reduction of Iron or Manganese. Applied and Environmental Microbiology 54:1472–1480.

7. Light SH, Méheust R, Ferrell JL, Cho J, Deng D, Agostoni M, Iavarone AT, Banfield JF, D’Orazio SEF, Portnoy DA. 2019. Extracellular electron transfer powers flavinylated extracellular reductases in Gram-positive bacteria. Proceedings of the National Academy of Sciences 116:26892–26899.

8. Glasser NR, Kern SE, Newman DK. 2014. Phenazine redox cycling enhances anaerobic survival in Pseudomonas aeruginosa by facilitating generation of ATP and a proton-motive force. Mol Microbiol 92:399–412.

9. Ciemniecki JA, Ho C-L, Horak RD, Okamoto A, Newman DK. 2024. Mechanistic study of a low-power bacterial maintenance state using high-throughput electrochemistry. Cell 187:6882–6895.e8.

10. Kundu BB, Krishnan J, Szubin R, Patel A, Palsson BO, Zielinski DC, Ajo-Franklin CM. 2025. Extracellular respiration is a latent energy metabolism in Escherichia coli. Cell 188:2907–2924.e23.

11. Lovley DR, Holmes DE. 2022. Electromicrobiology: the ecophysiology of phylogenetically diverse electroactive microorganisms. Nat Rev Microbiol 20:5–19.

12. Bretschger O, Obraztsova A, Sturm CA, Chang IS, Gorby YA, Reed SB, Culley DE, Reardon CL, Barua S, Romine MF, Zhou J, Beliaev AS, Bouhenni R, Saffarini D, Mansfeld F, Kim B-H, Fredrickson JK, Nealson KH. 2007. Current Production and Metal Oxide Reduction by Shewanella oneidensis MR-1 Wild Type and Mutants. Applied and Environmental Microbiology 73:7003–7012.

13. Li S, Zuo X, Carpenter MD, Verduzco R, Ajo-Franklin CM. 2025. Microbial bioelectronic sensors for environmental monitoring. Nat Rev Bioeng 3:30–49.

14. Chen H, Dong F, Minteer SD. 2020. The progress and outlook of bioelectrocatalysis for the production of chemicals, fuels and materials. Nat Catal 3:225–244.

15. Wang X, Aulenta F, Puig S, Esteve-Núñez A, He Y, Mu Y, Rabaey K. 2020. Microbial electrochemistry for bioremediation. Environmental Science and Ecotechnology 1:100013.

16. Nealson KH. 2017. Bioelectricity (electromicrobiology) and sustainability. Microb Biotechnol 10:1114–1119.

17. Graham AJ, Keitz BK. 2022. Living Synthetic Polymerizations, p. 27–49. In Srubar III, WV (ed.), Engineered Living Materials. Springer International Publishing, Cham.

18. Conley BE, Weinstock MT, Bond DR, Gralnick JA. 2020. A Hybrid Extracellular Electron Transfer Pathway Enhances the Survival of Vibrio natriegens. Applied and Environmental Microbiology 86:e01253–20.

19. Gong Z, Xie R, Zhang Y, Wang M, Tan T. 2023. Identification of Emerging Industrial Biotechnology Chassis Vibrio natriegens as a Novel High Salt-Tolerant and Feedstock Flexibility Electroactive Microorganism for Microbial Fuel Cell. Microorganisms 11:490.

20. Hoffart E, Grenz S, Lange J, Nitschel R, Müller F, Schwentner A, Feith A, Lenfers-Lücker M, Takors R, Blombach B. 2017. High Substrate Uptake Rates Empower Vibrio natriegens as Production Host for Industrial Biotechnology. Applied and Environmental Microbiology 83:e01614–17.

21. Eagon RG. 1962. Pseudomonas natriegens, a marine bacterium with a generation time of less than 10 minutes. Journal of Bacteriology 83:736–737.

22. Lima M, Muddana C, Xiao Z, Bandyopadhyay A, Wangikar PP, Pakrasi HB, Tang YJ. 2024. The new chassis in the flask: Advances in *Vibrio natriegens* biotechnology research. Biotechnology Advances 77:108464.

23. Gemünde A, Gail J, Holtmann D. 2023. Anodic Respiration of Vibrio natriegens in a Bioelectrochemical System. ChemSusChem 16:e202300181.

24. Hoff J, Daniel B, Stukenberg D, Thuronyi BW, Waldminghaus T, Fritz G. 2020. Vibrio natriegens: an ultrafast-growing marine bacterium as emerging synthetic biology chassis. Environmental Microbiology 22:4394–4408.

25. Su C, Cui H, Wang W, Liu Y, Cheng Z, Wang C, Yang M, Qu L, Li Y, Cai Y, He S, Zheng J, Zhao P, Xu P, Dai J, Tang H. 2025. Bioremediation of complex organic pollutants by engineered Vibrio natriegens. Nature 642:1024–1033.

26. Smith M, Hernández JS, Messing S, Ramakrishnan N, Higgins B, Mehalko J, Perkins S, Wall VE, Grose C, Frank PH, Cregger J, Le PV, Johnson A, Sherekar M, Pagonis M, Drew M, Hong M, Widmeyer SRT, Denson J-P, Snead K, Poon I, Waybright T, Champagne A, Esposito D, Jones J, Taylor T, Gillette W. 2024. Producing recombinant proteins in Vibrio natriegens. Microbial Cell Factories 23:208.

27. Hädrich M, Scheuchenegger C, Vital S-T, Gunkel C, Müller S, Hoff J, Borger J, Glawischnig E, Thoma F, Blombach B. 2025. Low-biomass pyruvate production with engineered Vibrio natriegens is accompanied by parapyruvate formation. Microbial Cell Factories 24:73.

28. Baker IR, Conley BE, Gralnick JA, Girguis PR. 2022. Evidence for Horizontal and Vertical Transmission of Mtr-Mediated Extracellular Electron Transfer among the Bacteria. mBio 13:e02904–21.

29. Gralnick JA, Bond DR. 2023. Electron Transfer Beyond the Outer Membrane: Putting Electrons to Rest. Annual Review of Microbiology 77:517–539.

30. Conley BE, Intile PJ, Bond DR, Gralnick JA. 2018. Divergent Nrf Family Proteins and MtrCAB Homologs Facilitate Extracellular Electron Transfer in Aeromonas hydrophila. Appl Environ Microbiol 84:e02134–18.

31. Coursolle D, Gralnick JA. 2012. Reconstruction of Extracellular Respiratory Pathways for Iron(III) Reduction in Shewanella Oneidensis Strain MR-1. Front Microbiol 3.

32. Reardon CL, Dohnalkova AC, Nachimuthu P, Kennedy DW, Saffarini DA, Arey BW, Shi L, Wang Z, Moore D, Mclean JS, Moyles D, Marshall MJ, Zachara JM, Fredrickson JK, Beliaev AS. 2010. Role of outer-membrane cytochromes MtrC and OmcA in the biomineralization of ferrihydrite by Shewanella oneidensis MR-1. Geobiology 8:56–68.

33. Marshall MJ, Beliaev AS, Dohnalkova AC, Kennedy DW, Shi L, Wang Z, Boyanov MI, Lai B, Kemner KM, McLean JS, Reed SB, Culley DE, Bailey VL, Simonson CJ, Saffarini DA, Romine MF, Zachara JM, Fredrickson JK. 2006. c-Type cytochrome-dependent formation of U(IV) nanoparticles by Shewanella oneidensis. PLoS Biol 4:e268.

34. Sadrzadeh SM, Graf E, Panter SS, Hallaway PE, Eaton JW. 1984. Hemoglobin. A biologic fenton reagent. Journal of Biological Chemistry 259:14354–14356.

35. Grove J, Tanapongpipat S, Thomas G, Griffiths L, Crooke H, Cole J. 1996. Escherichia coli K-12 genes essential for the synthesis of c-type cytochromes and a third nitrate reductase located in the periplasm. Molecular Microbiology 19:467–481.

36. Fuchs H, Ullrich SR, Hedrich S. 2024. Vibrio natriegens as a superior host for the production of c-type cytochromes and difficult-to-express redox proteins. Sci Rep 14:6093.

37. Goldbeck CP, Jensen HM, TerAvest MA, Beedle N, Appling Y, Hepler M, Cambray G, Mutalik V, Angenent LT, Ajo-Franklin CM. 2013. Tuning Promoter Strengths for Improved Synthesis and Function of Electron Conduits in Escherichia coli. ACS Synth Biol 2:150–159.

38. Su L, Fukushima T, Prior A, Baruch M, Zajdel TJ, Ajo-Franklin CM. 2020. Modifying Cytochrome c Maturation Can Increase the Bioelectronic Performance of Engineered Escherichia coli. ACS Synth Biol 9:115–124.

39. Park Y, Espah Borujeni A, Gorochowski TE, Shin J, Voigt CA. 2020. Precision design of stable genetic circuits carried in highly-insulated E. coli genomic landing pads. Mol Syst Biol 16:e9584.

40. Meyer AJ, Segall-Shapiro TH, Glassey E, Zhang J, Voigt CA. 2019. Escherichia coli “Marionette” strains with 12 highly optimized small-molecule sensors. Nat Chem Biol 15:196–204.

41. Sun Y, Xu J, Zhou H, Zhang H, Wu J, Yang L. 2023. Recombinant Protein Expression Chassis Library of Vibrio natriegens by Fine-Tuning the Expression of T7 RNA Polymerase. ACS Synth Biol 12:555–564.

42. Kranz RG, Sutherland MC. 2025. Mechanisms and Control of Heme Transport and Incorporation into Cytochrome c. Annual Review of Microbiology 79:23–43.

43. Schicklberger M, Bücking C, Schuetz B, Heide H, Gescher J. 2011. Involvement of the Shewanella oneidensis Decaheme Cytochrome MtrA in the Periplasmic Stability of the β-Barrel Protein MtrB. Applied and Environmental Microbiology 77:1520–1523.

44. Stukenberg D, Hoff J, Faber A, Becker A. 2022. NT-CRISPR, combining natural transformation and CRISPR-Cas9 counterselection for markerless and scarless genome editing in Vibrio natriegens. Commun Biol 5:265.

45. Wu F, Chen W, Peng Y, Tu R, Lin Y, Xing J, Wang Q. 2020. Design and Reconstruction of Regulatory Parts for Fast-Growing Vibrio natriegens Synthetic Biology. ACS Synth Biol 9:2399–2409.

46. Bhakta SP. 2024. A Bacterial Toolkit for Rapid Prototyping of Multicistronic Genetic Circuits from Interchangeable Parts. Rice University.

47. Chang H-Y, Ahn Y, Pace LA, Lin MT, Lin Y-H, Gennis RB. 2010. The Diheme Cytochrome c4 from Vibrio cholerae Is a Natural Electron Donor to the Respiratory cbb3 Oxygen Reductase. Biochemistry 49:7494–7503.

48. Campbell IJ, Atkinson JT, Carpenter MD, Myerscough D, Su L, Ajo-Franklin CM, Silberg JJ. 2022. Determinants of Multiheme Cytochrome Extracellular Electron Transfer Uncovered by Systematic Peptide Insertion. Biochemistry 61:1337–1350.

49. Coursolle D, Gralnick JA. 2010. Modularity of the Mtr respiratory pathway of Shewanella oneidensis strain MR-1. Molecular Microbiology 77:995–1008.

50. Bücking C, Piepenbrock A, Kappler A, Gescher J. 2012. Outer-membrane cytochrome-independent reduction of extracellular electron acceptors in Shewanella oneidensis. Microbiology 158:2144–2157.

51. Marsili E, Baron DB, Shikhare ID, Coursolle D, Gralnick JA, Bond DR. 2008. Shewanella secretes flavins that mediate extracellular electron transfer. Proceedings of the National Academy of Sciences 105:3968–3973.

52. Mevers E, Su L, Pishchany G, Baruch M, Cornejo J, Hobert E, Dimise E, Ajo-Franklin CM, Clardy J. 2019. An elusive electron shuttle from a facultative anaerobe. eLife 8:e48054.

53. Oram J, Jeuken LJC. 2016. A Re-evaluation of Electron-Transfer Mechanisms in Microbial Electrochemistry: Shewanella Releases Iron that Mediates Extracellular Electron Transfer. ChemElectroChem 3:829–835.

54. Abuyen K, El-Naggar MY. 2023. Soluble Iron Enhances Extracellular Electron Uptake by Shewanella oneidensis MR-1. ChemElectroChem 10:e202200965.

55. Delgado VP, Paquete CM, Sturm G, Gescher J. 2019. Improvement of the electron transfer rate in Shewanella oneidensis MR-1 using a tailored periplasmic protein composition. Bioelectrochemistry 129:18–25.

56. Sun W, Lin Z, Yu Q, Cheng S, Gao H. 2021. Promoting Extracellular Electron Transfer of Shewanella oneidensis MR-1 by Optimizing the Periplasmic Cytochrome c Network. Front Microbiol 12.

57. Stevens DM, Tang A, Coaker G. 2021. A Genetic Toolkit for Investigating Clavibacter Species: Markerless Deletion, Permissive Site Identification, and an Integrative Plasmid. Mol Plant Microbe Interact 34:1336–1345.

58. Zwietering MH, Jongenburger I, Rombouts FM, van ’t Riet K. 1990. Modeling of the Bacterial Growth Curve. Applied and Environmental Microbiology 56:1875–1881.

59. Lew M. 2007. Good statistical practice in pharmacology. Problem 2. Br J Pharmacol 152:299–303.

60. Zweifach A. 2024. Samples in many cell-based experiments are matched/paired but taking this into account does not always increase power of statistical tests for differences in means. MBoC 35:br1.

