## Supplemental Information for "Electrode Reduction by *Vibrio natriegens* Depends on Balanced Expression of Multiheme Cytochromes"

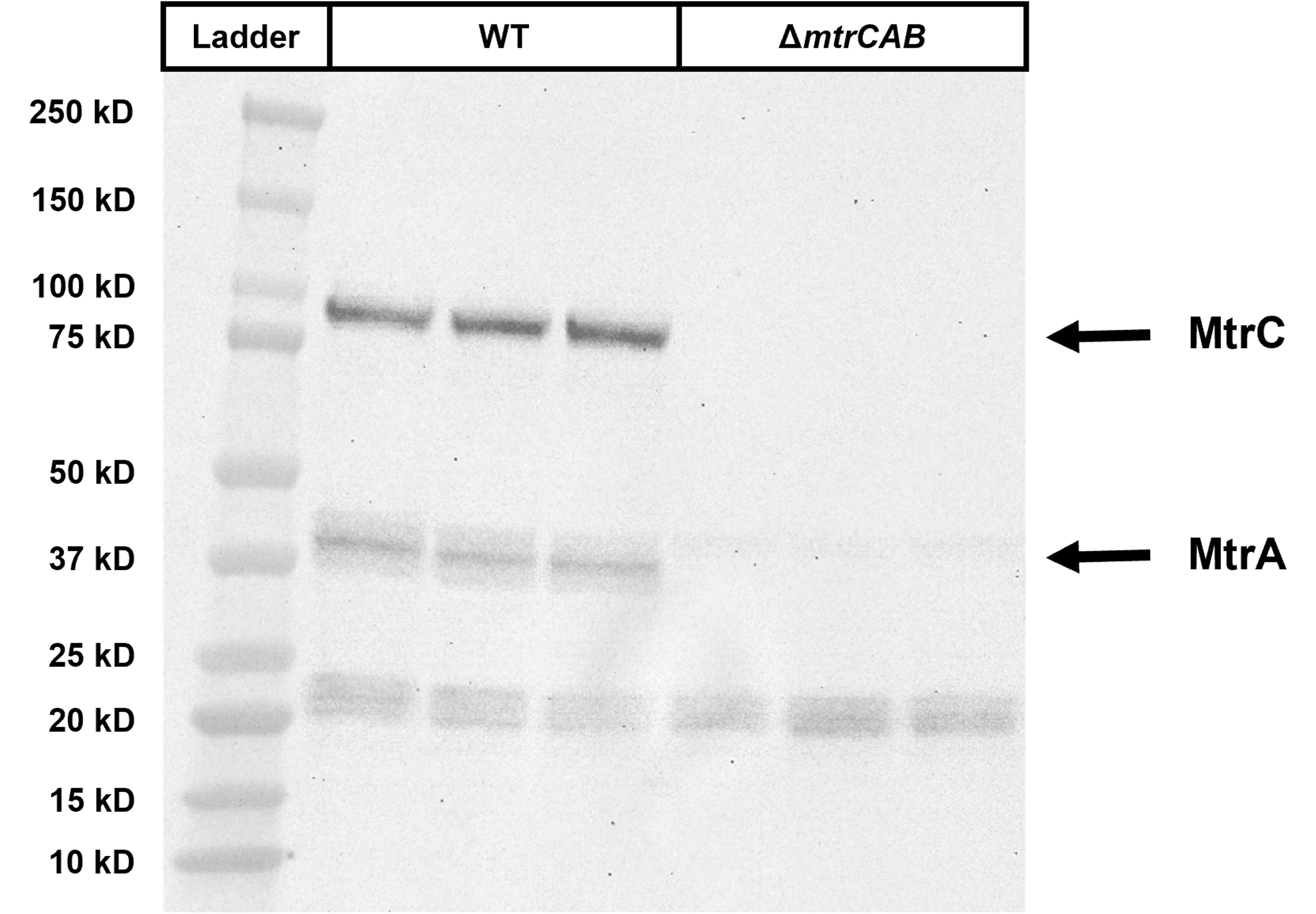

Supplemental Figure 1: Aerobic cultivation of *V. natriegens* in rich medium supports production of multiheme cytochromes *c*. Comparison of the band patterns produced by ECL of whole cell lysates of WT and Δ*mtrCAB* *V. natriegens* (three biological replicates each) shows that WT cells produce bands at the expected molecular weights for MtrA and MtrC that are absent in cells lacking the genes for the Mtr porin:cytochrome *c* complex.

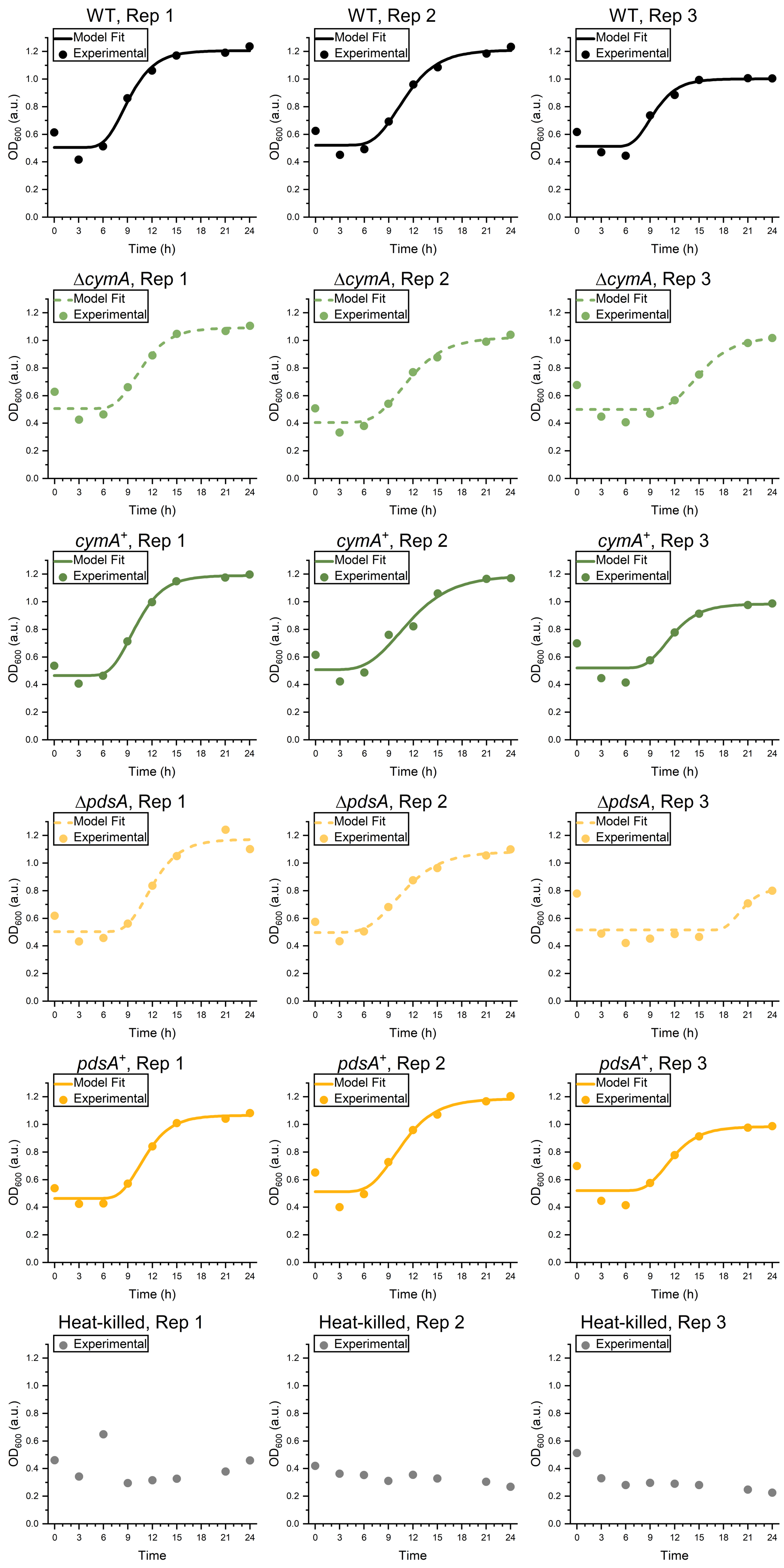

Supplemental Figure 2: Growth curves show that live cells of all *cymA* and *pdsA* genotypes grow inside BESs. Experimental measurements (circles) of OD_600_ sampled from BESs are plotted alongside functions (lines) produced by fitting the Zwietering-modified Gompertz equation for microbial growth to the experimental data (functions were not fitted to the data for heat-killed cells). All measurements were made concurrently with the current density measurements in **Figure 2**. Each row contains three independent biological replicates for a given strain. Each column represents a block of samples that were measured on the same day.

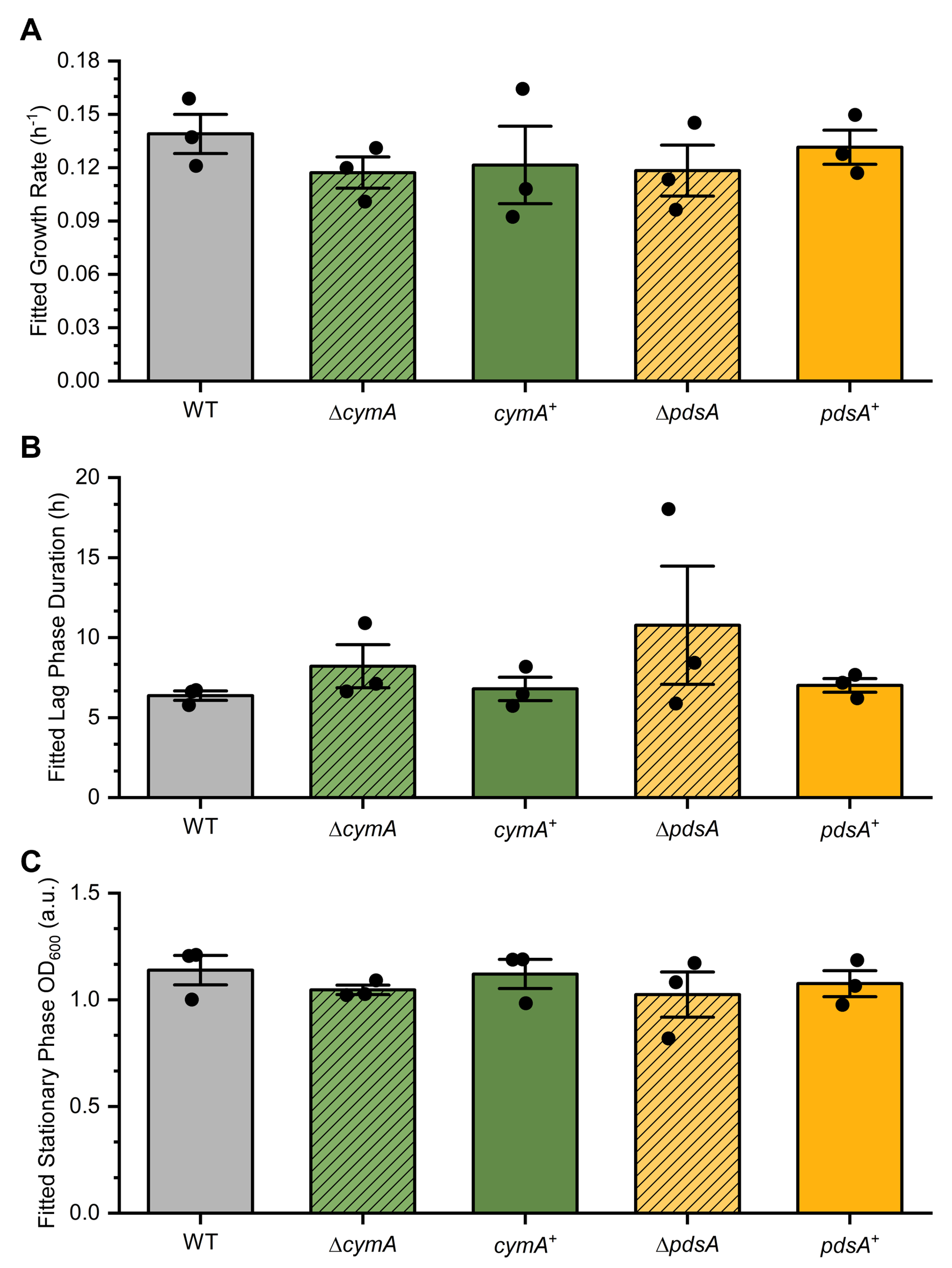

Supplemental Figure 3: Growth curve parameters do not significantly differ among different *cymA* and *pdsA* genotypes. Parameters corresponding to growth rate **(A)**, duration of lag phase **(B)**, and stationary phase OD_600_ **(C)** were extracted from the Zwietering-modified Gompertz models fitted in **Supplemental Figure 2**. Random block ANOVA did not find any significant differences among the tested strains for the growth rate (p = 0.2410), duration of lag phase (p = 0.3191), or stationary phase OD_600_ (p = 0.3250) parameters.

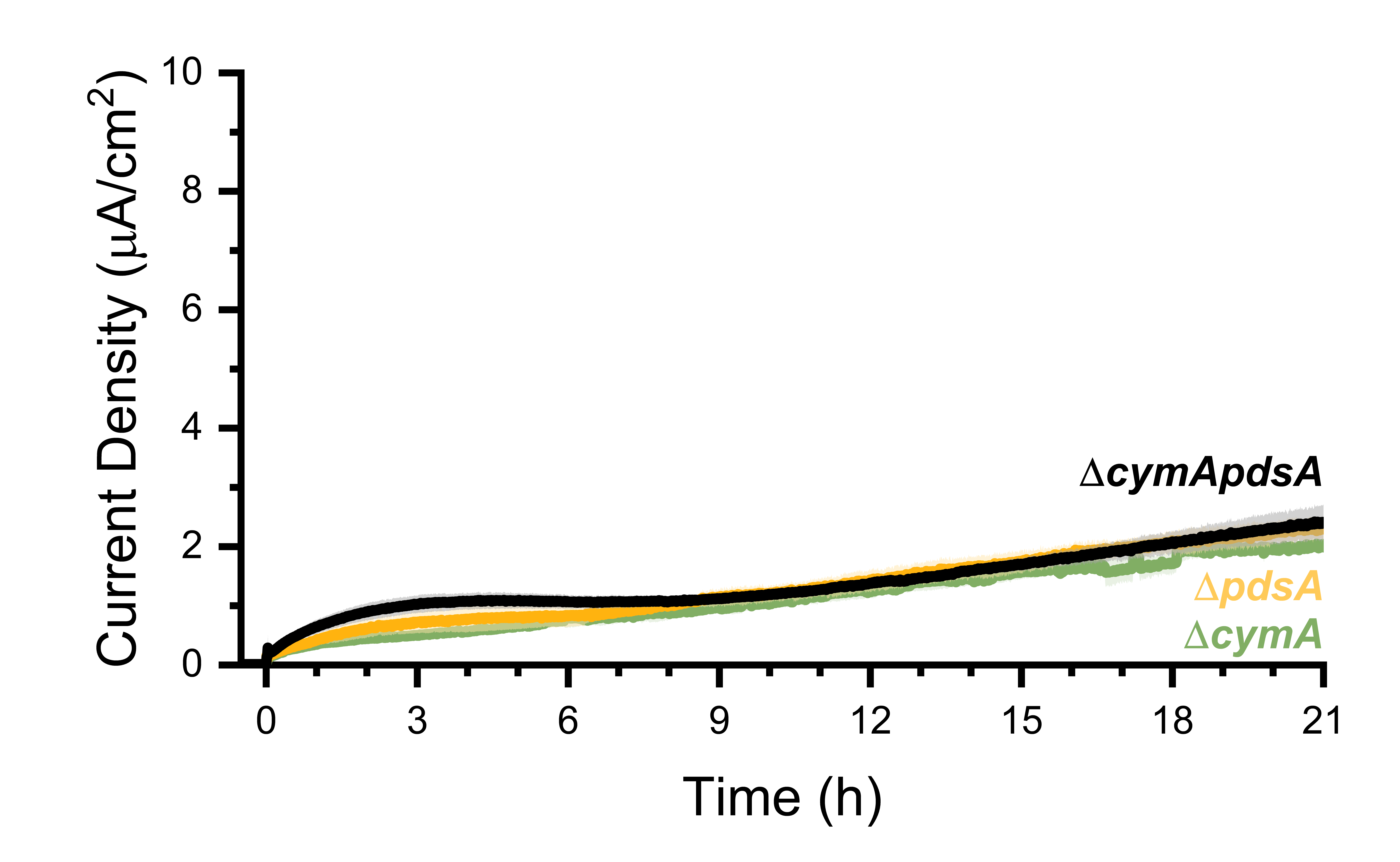

Supplemental Figure 4: Δ*cymApdsA* *V. natriegens* produces similar current to Δ*cymA* and Δ*pdsA*. Chronoamperometry of Δ*cymA* (light green) and Δ*pdsA* (light gold) single knock-out strains compared to a Δ*cymApdsA* (black) double knock-out strain shows no discernible difference. Each curve represents the mean of three biological replicates. Error bars indicate standard error of the mean.

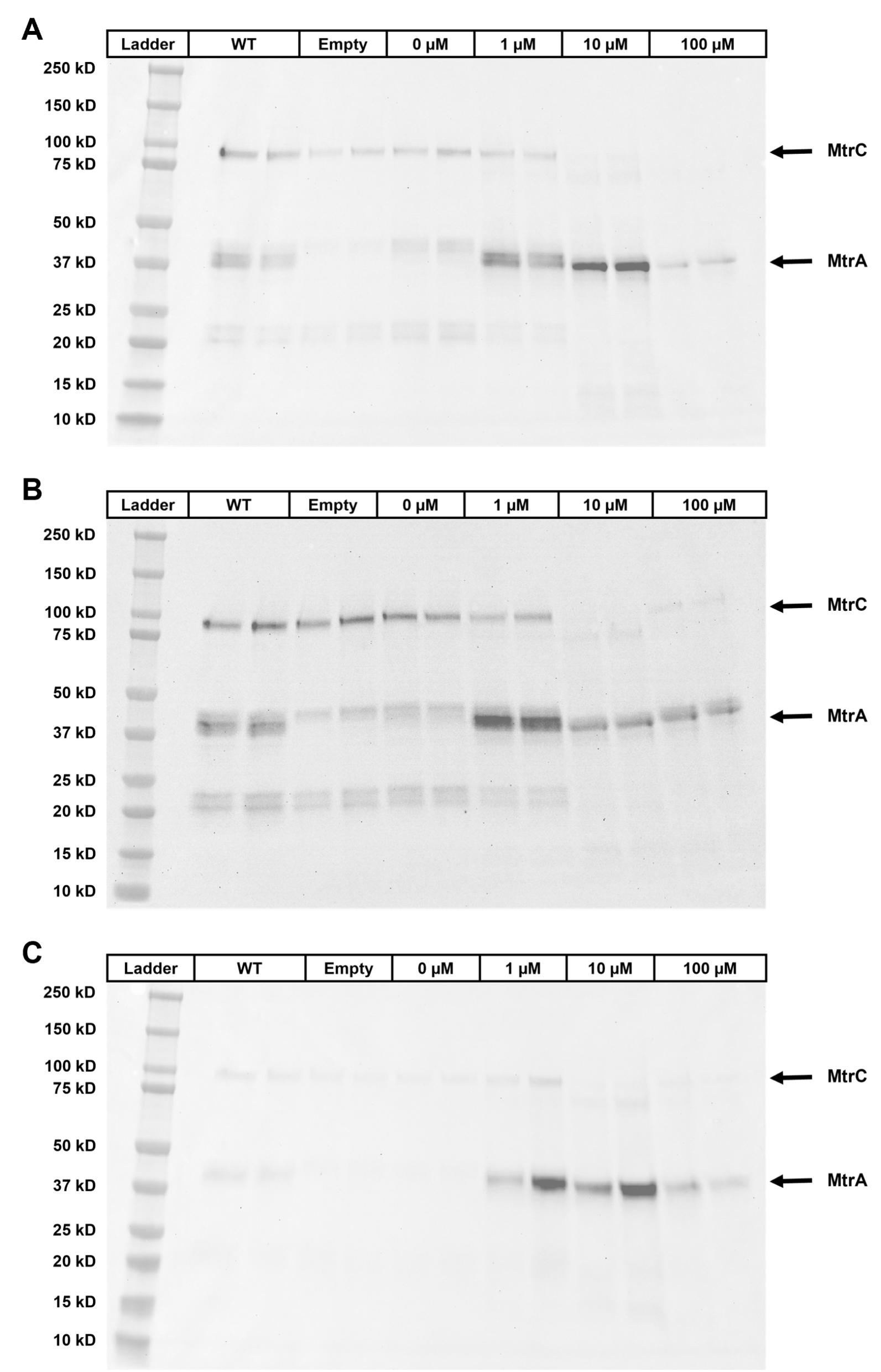

Supplemental Figure 5: Induction of MtrA expression in *V. natriegens* alters the balance of cytochromes**.** ECL of whole cell lysates of cell suspensions immediately prior to injection into BESs in **Figure 4A**. Each cell suspension was lysed and loaded in technical duplicate. Panels A, B, and C each correspond to a different experimental replicate, performed on separate days. The samples within each Panel correspond to BESs operated in parallel.

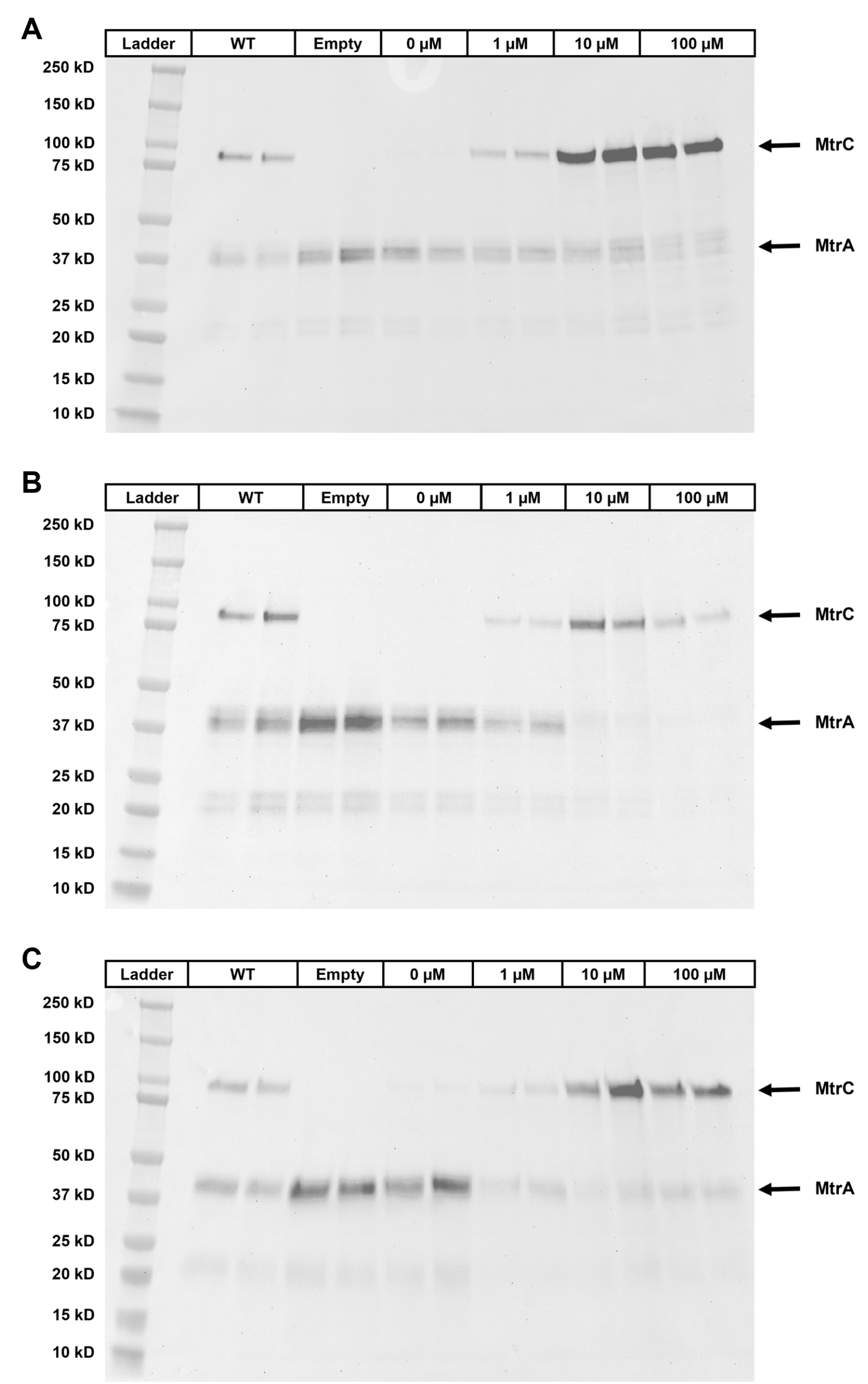

Supplemental Figure 6: Induction of MtrC expression in *V. natriegens* alters the balance of cytochromes**.** ECL of whole cell lysates of cell suspensions immediately prior to injection into BESs in **Figure 4B**. Each cell suspension was lysed and loaded in technical duplicate. Panels A, B, and C each correspond to a different experimental replicate, performed on separate days. The samples within each Panel correspond to BESs operated in parallel.

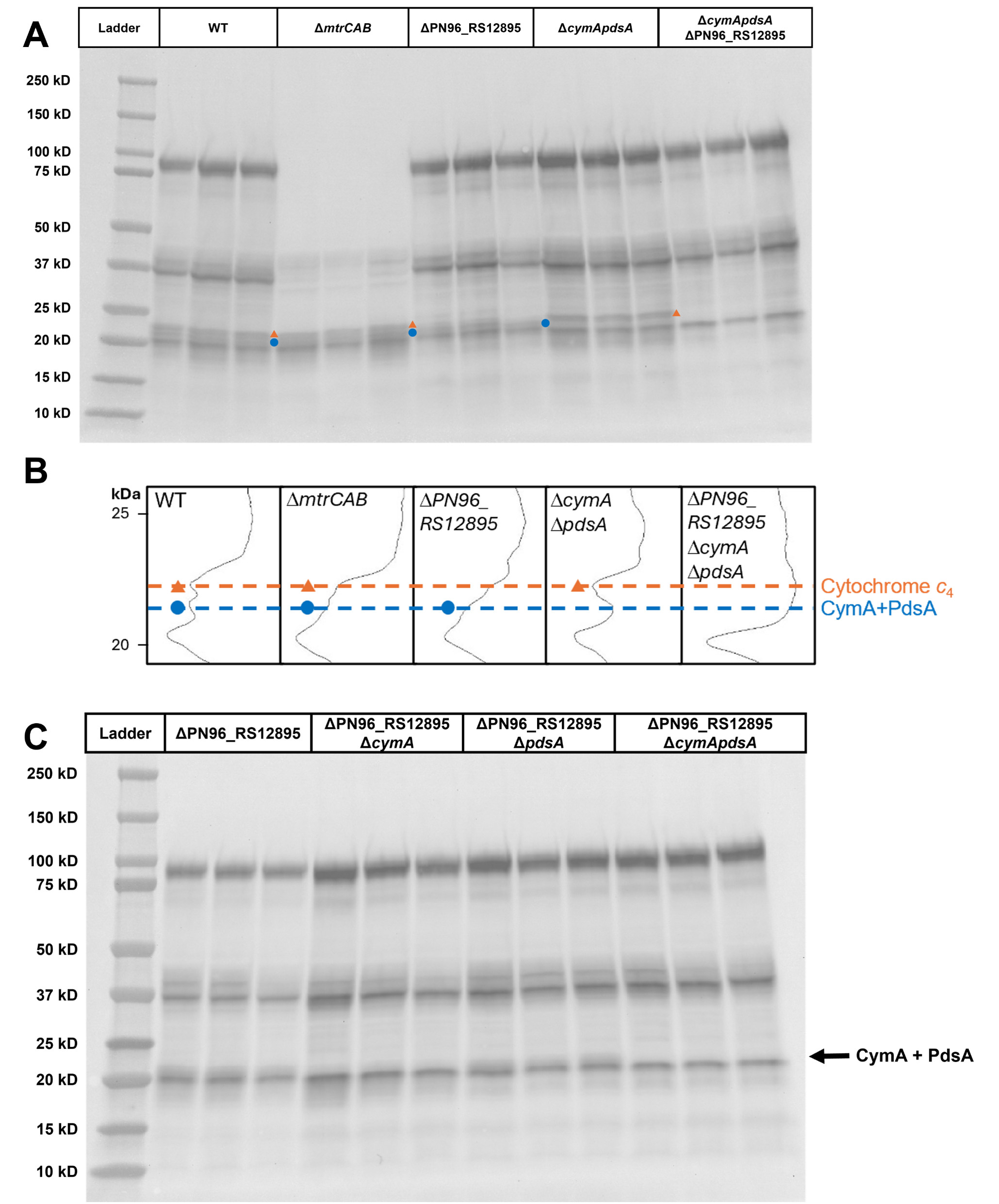

Supplemental Figure 7: Deletion of Cytochrome *c*_4_ facilitates identification of a band attributable to the combination of CymA and PdsA**. (A)** Comparison of the band patterns produced by ECL of whole cell lysates of WT, Δ*mtrCAB*, Δ*cymApdsA*, ΔPN96_RS12895, and Δ*cymApdsA*PN96_RS12895 (three biological replicates each) shows that the strains carrying *cymA* and *pdsA* produce a band (blue circle) in the range 20-25 kDa that is more easily visualized when PN96_RS12895 (orange triangle) is deleted. **(B)** Densitometry of representative samples from **A**. **(C)** Comparison of the band patterns produced by ECL of whole cell lysates of ΔPN96_RS12895, Δ*cymA*PN96_RS12895, Δ*pdsA*PN96_RS12895, and Δ*cymApdsA*PN96_RS12895 (three biological replicates each) shows that both *cymA* and *pdsA* contribute to a band (labeled) that is eliminated when both genes are deleted.

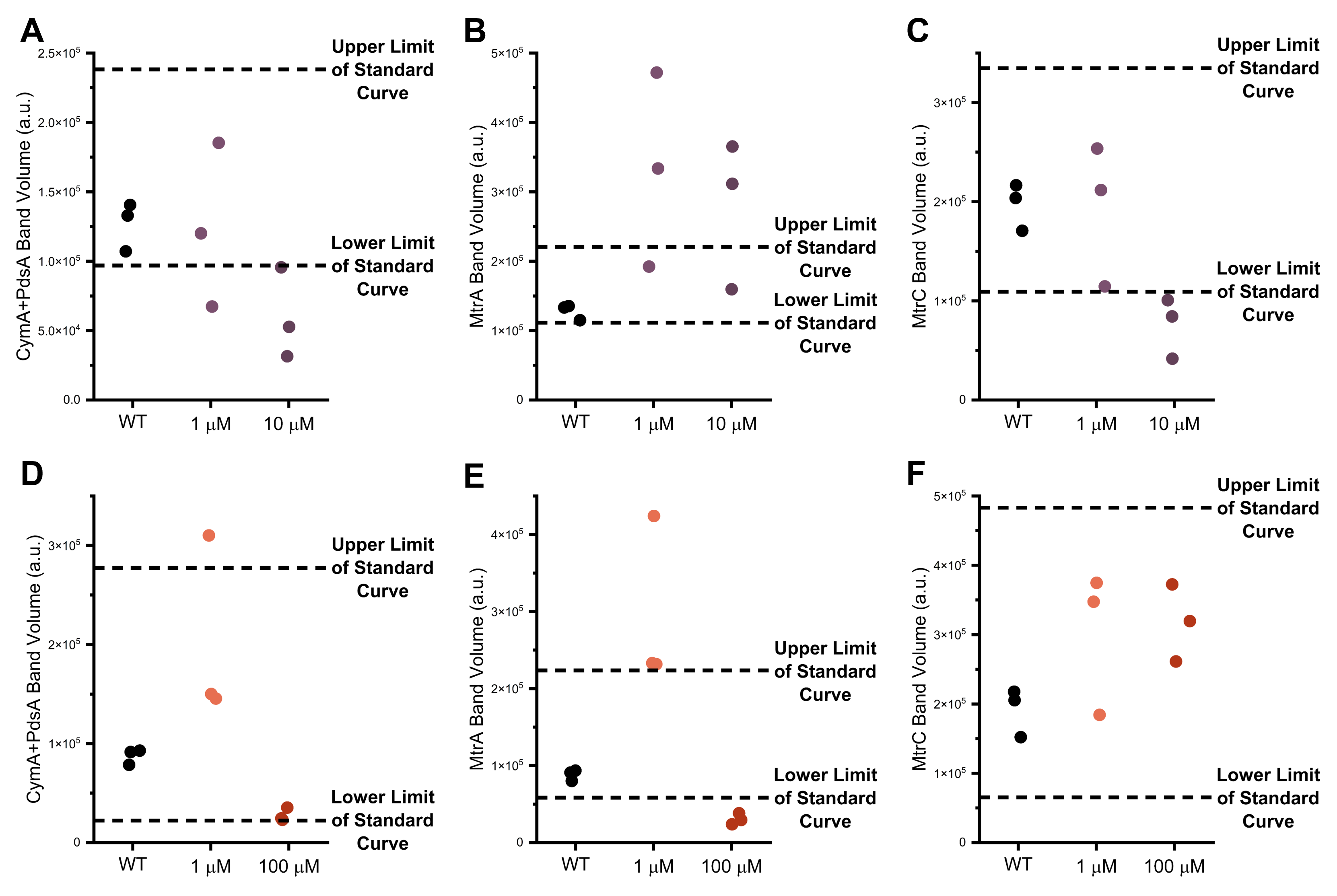

Supplemental Figure 8: Image analysis of cytochrome *c* band patterns finds that overexpression of either *mtrA* or *mtrC* yields lower band volumes for other cytochromes**.** The ECL visualized blots from **Figure 5** were used to extract band volumes for cytochromes of interest. **(A)** Comparison of the CymA+PdsA band volume across *mtrA* cumate induction conditions shows that 10 µM cumate-treated cells produced lower band volumes than WT cells. **(B)** Comparison of the MtrA band volume across *mtrA* cumate induction conditions shows that cumate induction resulted in band volumes greater than those observed from WT cells. **(C)** Comparison of the MtrC band volume across *mtrA* cumate induction conditions shows that 10 µM cumate-treated cells produced lower band volumes than WT cells. **(D)** Comparison of the CymA+PdsA band volume across *mtrC* cumate induction conditions shows that 100 µM cumate-treated cells produced lower band volumes than WT cells. **(E)** Comparison of the MtrA band volume across *mtrC* cumate induction conditions shows that 100 µM cumate-treated cells produced lower band volumes than WT cells. **(F)** Comparison of the MtrC band volume across *mtrC* cumate induction conditions shows that cumate induction resulted in band volumes similar to or greater than those observed from WT cells. **Figure 5A,B** features a standard curve of WT cells loaded at five different concentrations. The dashed lines represent the band volumes of the indicated band extracted from the lowest and highest concentration standard curve lanes. The three data points for each condition represent three biological replicates.

**Supplemental Table 1**

| **Strain** | **Chromosomal Modification(s)** | **Plasmid** | **Source** |
| --- | --- | --- | --- |
| *Vibrio natriegens* ATCC 14048 | None | None | Gift from George Bennett, Rice University |
| MDC069 | None | pMC53 | This study; transformation of *V. natriegens* ATCC 14048 with pMC53 |
| MDC070 | None | pMC54 | This study; transformation of *V. natriegens* ATCC 14048 with pMC54 |
| MDC071 | Δ*cymA* | pMC53 | This study; natural transformation of MDC069 with tDNA_cymA |
| MDC072 | Δ*cymA* | None | This study; curing pMC53 from MDC071 |
| MDC076 | Δ*pdsA* | pMC54 | This study; natural transformation of MDC070 with tDNA_pdsA |
| MDC077 | Δ*pdsA* | None | This study; curing pMC54 from MDC076 |
| MDC081 | None | pMC67 | This study; transformation of *V. natriegens* ATCC 14048 with pMC67 |
| MDC082 | Δ*mtrC* | pMC67 | This study; natural transformation of MDC081 with tDNA_mtrC |
| MDC083 | Δ*mtrCAB* | pMC67 | This study; natural transformation of MDC081 with tDNA_mtrCAB |
| MDC084 | Δ*mtrC* | None | This study; curing pMC67 from MDC082 |
| MDC085 | Δ*mtrCAB* | None | This study; curing pMC67 from MDC083 |
| MDC089 | None | pMC69 | This study; transformation of *V. natriegens* ATCC 14048 with pMC69 |
| MDC091 | Δ*mtrB* | pMC69 | This study; natural transformation of MDC089 with tDNA_mtrB |
| MDC092 | Δ*mtrA* | pMC70 | This study; natural transformation of WCC36 with tDNA_mtrA |
| MDC093 | Δ*mtrA* | None | This study; curing pMC70 from MDC092 |
| MDC094 | Δ*mtrB* | None | This study; curing pMC69 from MDC091 |
| MDC121 | Δ*mtrA* | pWC12 | This study; transformation of MDC093 with pWC12 |
| MDC122 | Δ*mtrC* | pWC12 | This study; transformation of MDC084 with pWC12 |
| MDC123 | Δ*mtrA*;  LP2::*cymR* | pWC12 | This study; natural transformation of MDC121 with tDNA_cymR |
| MDC124 | Δ*mtrC*;  LP2::*cymR* | pWC12 | This study; natural transformation of MDC122 with tDNA_cymR |
| MDC125 | Δ*mtrA*;  LP2::*cymR* | None | This study; curing pWC12 from MDC123 |
| MDC126 | Δ*mtrC*;  LP2::*cymR* | None | This study; curing pWC12 from MDC124 |
| MDC127 | Δ*mtrA*;  LP2::*cymR* | pWC1 | This study; transformation of MDC125 with pWC1 |
| MDC128 | Δ*mtrC*;  LP2::*cymR* | pWC1 | This study; transformation of MDC126 with pWC1 |
| MDC135 | Δ*mtrC*;  LP1::P_cymRD_-*mtrC*; LP2::*cymR* | None | This study; curing pWC1 from MDC141 |
| MDC139 | Δ*mtrA*;  LP1::P_cymRD_-*mtrA*; LP2::*cymR* | pWC1 | This study; natural transformation of MDC127 with tDNA_PcymRD-mtrA |
| MDC141 | Δ*mtrC*;  LP1::P_cymRD_-*mtrC*; LP2::*cymR* | pWC1 | This study; natural transformation of MDC128 with tDNA_PcymRD-mtrC |
| MDC142 | Δ*mtrA*;  LP1::P_cymRD_-*mtrA*; LP2::*cymR* | None | This study; curing pWC1 from MDC139 |
| MDC153 | Δ*mtrA* | pWC24 | This study; transformation of MDC093 with pWC24 |
| MDC154 | Δ*mtrC* | pWC24 | This study; transformation of MDC084 with pWC24 |
| MDC155 | Δ*mtrA*;  LP1::P_cymRD_-*mtrA*; LP2::*cymR* | pWC24 | This study; transformation of MDC142 with pWC24 |
| MDC156 | Δ*mtrC*;  LP1::P_cymRD_-*mtrC*; LP2::*cymR* | pWC24 | This study; transformation of MDC135 with pWC24 |
| MDC159 | Δ*mtrAPN96_RS12895* | pWC24 | This study; natural transformation of MDC153 with tDNA_PN96_RS12895 |
| MDC160 | Δ*mtrCPN96_RS12895* | pWC24 | This study; natural transformation of MDC154 with tDNA_PN96_RS12895 |
| MDC161 | Δ*mtrAPN96_RS12895*;  LP1::P_cymRD_-*mtrA*; LP2::*cymR* | pWC24 | This study; natural transformation of MDC155 with tDNA_PN96_RS12895 |
| MDC162 | Δ*mtrCPN96_RS12895*;  LP1::P_cymRD_-*mtrC*; LP2::*cymR* | pWC24 | This study; natural transformation of MDC156 with tDNA_PN96_RS12895 |
| MDC165 | Δ*mtrAPN96_RS12895* | None | This study; curing pWC24 from MDC159 |
| MDC166 | Δ*mtrCPN96_RS12895* | None | This study; curing pWC24 from MDC160 |
| MDC167 | Δ*mtrAPN96_RS12895*;  LP1::P_cymRD_-*mtrA*; LP2::*cymR* | None | This study; curing pWC24 from MDC161 |
| MDC168 | Δ*mtrCPN96_RS12895*;  LP1::P_cymRD_-*mtrC*; LP2::*cymR* | None | This study; curing pWC24 from MDC162 |
| WCC1 | Δ*cymA* | pWC1 | This study; transformation of MDC072 with pWC1 |
| WCC2 | Δ*pdsA* | pWC1 | This study; transformation of MDC077 with pWC1 |
| WCC15 | Δ*cymA*;  LP1::P_cymRC_-*cymA* | pWC1 | This study; natural transformation of WCC1 with tDNA_PcymRC-cymA |
| WCC19 | Δ*pdsA*;  LP1::P_cymRC_-*pdsA* | pWC1 | This study; natural transformation of WCC2 with tDNA_PcymRC-pdsA |
| WCC24 | Δ*cymA*;  LP1::P_cymRC_-*cymA* | None | This study; curing pWC1 from WCC15 |
| WCC28 | Δ*pdsA*;  LP1::P_cymRC_-*pdsA* | None | This study; curing pWC1 from WCC19 |
| WCC36 | None | pMC70 | This study; transformation of *V*. *natriegens* ATCC 14048 with pMC70 |
| WCC50 | Δ*mtrB* | pWC1 | This study; transformation of MDC094 with pWC1 |
| WCC53 | Δ*mtrB*;  LP1::P_cymRC_-*mtrB* | None | This study; curing pWC1 from WCC57 |
| WCC57 | Δ*mtrB*;  LP1::P_cymRC_-*mtrB* | pWC1 | This study; natural transformation of WCC50 with tDNA_PcymRC-mtrB |
| WCC70 | Δ*pdsA* | pMC53 | This study; transformation of MDC077 with pMC53 |
| WCC71 | Δ*cymApdsA* | pMC53 | This study; natural transformation of WCC70 with tDNA_cymA_2 |
| WCC72 | Δ*cymApdsA* | None | This study; curing pMC53 from WCC71 |
| WCC83 | None | pWC24 | This study; transformation of *V. natriegens* ATCC 14048 with pWC24 |
| WCC86 | Δ*cymApdsA* | pWC24 | This study; transformation of WCC72 with pWC24 |
| WCC89 | Δ*PN96_RS12895* | pWC24 | This study; natural transformation of WCC83 with tDNA_PN96_RS12895 |
| WCC92 | Δ*cymApdsAPN96_RS12895* | pWC24 | This study; natural transformation of WCC86 with tDNA_PN96_RS12895 |
| WCC94 | Δ*cymApdsAPN96_RS12895* | None | This study; curing pWC24 from WCC92 |
| WCC98 | Δ*PN96_RS12895* | None | This study; curing pWC24 from WCC89 |
| WCC99 | Δ*mtrC*;  LP1::P_cymRD_-Empty;  LP2::*cymR* | pWC1 | This study; natural transformation of MDC128 with tDNA_PcymRD-Empty |
| WCC100 | Δ*mtrA*;  LP1::P_cymRD_-Empty;  LP2::*cymR* | pWC1 | This study; natural transformation of MDC127 with tDNA_PcymRD-Empty |
| WCC101 | Δ*mtrC*;  LP1::P_cymRD_-Empty;  LP2::*cymR* | None | This study; curing pWC1 from WCC99 |
| WCC102 | Δ*mtrA*;  LP1::P_cymRD_-Empty;  LP2::*cymR* | None | This study; curing pWC1 from WCC100 |

Supplemental Table 1: List of *V. natriegens* strains**.** The highlighted strains were used in Figures; other strains were created as intermediates during the creation of the highlighted strains. All deletions consist of the exact removal of the full coding sequence (CDS) of each of the listed genes, except for Δ*mtrCAB* and Δ*mtrA*. For Δ*mtrCAB*, all bases from the start of the *mtrC* CDS to the end of the *mtrB* CDS were removed. For Δ*mtrA*, only bases 1-947 of the *mtrA* CDS were removed, leaving the final 40 bp of *mtrA* present in the genome. The 40 bp cutoff was chosen because it was the longest stretch of bases that could be retained without including a possible START codon.

**Supplemental Table 2**

| **Genomic Modification** | **gRNA Sequence** | **Single-stranded Oligonucleotides** | **NT-CRISPR Plasmid** | **Primers Used for Sequence Verification of Genomic Modifications** |
| --- | --- | --- | --- | --- |
| Deletion of *mtrCAB* | aatccacgctgactcccatg | gtccaatccacgctgactcccatg;  aaaccatgggagtcagcgtggatt | pMC67 | MC371; MC373 |
| Deletion of *cymA* | cgatcgtatccatcccaatg | gtcccgatcgtatccatcccaatg;  aaaccattgggatggatacgatcg | pMC53 | MC335; MC336 |
| Integration at LP1 | gggctaatatcttaaatgtg | gtccgggctaatatcttaaatgtg;  aaaccacatttaagatattagccc | pWC1 | WC27; WC28 |
| Deletion of *pdsA* | acgctcgattaattctggag | gtccacgctcgattaattctggag;  aaacctccagaattaatcgagcgt | pMC54 | MC335; MC336 |
| Deletion of *mtrA* | gatggttgctgagttgcaag | gtccgatggttgctgagttgcaag;  aaaccttgcaactcagcaaccatc | pMC70 | MC339; MC390 |
| Integration at LP2 | agtcgcttatggcgtgaaag | gtccagtcgcttatggcgtgaaag; aaacctttcacgccataagcgact | pWC12 | WC68; WC69 |
| Deletion of *mtrC* | aatccacgctgactcccatg | gtccaatccacgctgactcccatg;  aaaccatgggagtcagcgtggatt | pMC67 | MC371; MC372 |
| Deletion of PN96_RS12895 | accacgctcagcatcaccag | gtccaccacgctcagcatcaccag;  aaacctggtgatgctgagcgtggt | pWC24 | WC93; WC94 |
| Deletion of *mtrB* | tcacaagctacatctgaagg | gtcctcacaagctacatctgaagg;  aaacccttcagatgtagcttgtga | pMC69 | MC389; MC373 |

Supplemental Table 2: List of gRNA sequences, oligonucleotide pairs used to encode the gRNAs, and plasmids used to express the gRNAs to facilitate CRISPR-Cas9 counter-selection for the listed chromosomal modifications**.** In the oligonucleotide sequences, the overhangs used for Golden Gate assembly into pST140 are underlined.

Description of Supplemental Table 3: Plasmid sequences**.** gRNA sequences in plasmids are underlined, when present. Supplemental Table 3 is available as a separate Excel file.

Description of Supplemental Table 4: List of tDNA sequences used to perform chromosomal deletions and insertions using natural transformation**.** The Composition column lists the 5’ → 3’ order of genetic parts (if any) located between the upstream and downstream homology arms. For each tDNA, the DNA sequences of the upstream homology arm, any integrated genetic parts, and the downstream arm are listed in separate columns. Supplemental Table 4 is available as a separate Excel file.

**Supplemental Table 5**

| **Part Name** | **Part Type** | **Source** | **Sequence** |
| --- | --- | --- | --- |
| DT54 | Double Terminator | Park, *et al.* (1) | ggaaacacagaaaaaagcccgcacctgacagtgcgggctttttttttcgaccaaaggctcggtaccaaattccagaaaagacacccgaaagggtgttttttcgttttggtcc |
| DT101 | Double Terminator | Park, *et al.* (1) | tctttaaaaagaaacctccgcattgcggaggtttcgccttttgatactctgtctgaagtaattcttgccgcagtgaaaaatggcgcccatcggcgccatttttttatgcttccattagaaagcaaaaagcctgctagaaagcaggcttttttgaatttggctcctctgac |
| DT42 | Double Terminator | Park, *et al.* (1) | agttaaccaaaaaggggggattttatctcccctttaatttttcctcgcagatagcaaaaaagcgcctttagggcgcttttttacattggtgg |
| DT5 | Double Terminator | Park, *et al.* (1) | tccggcaattaaaaaagcggctaaccacgccgctttttttacgtctgcactcggtaccaaattccagaaaagaggcctcccgaaaggggggccttttttcgttttggtcc |
| P_cymRC_ | Promoter | Meyer, *et al.* (2) | aacaaacagacaatctggtctgtttgtattatggaaaatttttctgtataatagattcaacaaacagacaatctggtctgtttgtattat |
| P_cymRD_ | Promoter | (3) | gaaaacaaacagacaatccggtctgtttgtatacaggaaaatttttctgtataatagattcaacaaacagacaatctggtctgtttgtattat |
| B0034 | RBS | Sun, *et al.* (4) | aaagaggagaaatactag |
| P17 | Promoter-RBS | Wu, *et al.* (5) | aagcttatttgaaaaaagtgtttgacagcttctgaaggttattctataatgaagtctcgagttatcgagattttcaggagctaaggaagctaat |
| *cymR* | CDS (repressor) | *V. natriegens* ATCC 14048 | atgagcccgaaacgtcgtacccaggcagaacgtgcaatggaaacccagggtaaactgattgcagcagcactgggtgttctgcgtgaaaaaggttatgcaggttttcgtattgcagatgttccgggtgcagccggtgttagccgtggtgcacagagccatcattttccgaccaaactggaactgctgctggcaacctttgaatggctgtatgagcagattaccgaacgtagccgtgcacgtctggcaaaactgaaaccggaagatgatgttattcagcagatgctggatgatgcagcagaattttttctggatgatgattttagcatcggcctggatctgattgttgcagcagatcgtgatccggcactgcgtgaaggtattcagcgtaccgttgaacgtaatcgttttgttgttgaagatatgtggctgggtgtgctggtgagccgtggtctgagccgtgatgatgccgaagatattctgtggctgatttttaacagcgttcgtggtctggtagttcgtagcctgtggcagaaagataaagaacgttttgaacgtgtgcgtaatagcaccctggaaattgcacgtgaacgttatgcaaaattcaaacgttga |
| *cymA* | CDS (cytochrome) | *V. natriegens* ATCC 14048 | atgggatggtggagtcgacctaataaaaaatggatgctcggcatcccgcttggtggtgtagtcgcgtttatcttaggcgtgggcgcactaggcgcttaccatggcgttatgaattattccaacaccaacgacttctgtttcggatgccacattgggatggatacgatcgtggaggaatatcaagcgtcaatacattacaacaactcaaaaggtgtgattgccgcgacatgcagtgactgtcatgttccgagagagtttattcccaaaatgatcgttaaaatcaccgcgacagcggatatcttccataaattgaaaggcgatattacgctggaaaactttgagtcagaacaccggccaaggcttgcgacaatcgtaacggaggagtttgtcgccaataagtccaagcaatgtaaatactgtcaccaggttgatcgcatggacttcgaagcgcaaggtcgtacgacggctcgtcgacaccaaatgatggaatcacgagatcaatcgtgtatcgattgtcatgctggtatagcgcacaagttaccagaggatgagccagaagaagtcgaagaagaaatggttgaaaccaccgctatggttaattga |
| *pdsA* | CDS (cytochrome) | *V. natriegens* ATCC 14048 | atgtgttcttctttgacaaaaataactcaagcgttgacgttgattacagcaaactttcttctggtttcgtcagtgcaagcatccgctccagaattaatcgagcgtttggaatgtgaggcatgccatggaccgggtggggtatcgacaaataccaatattccttctattgcgggcttacctgaatttaatttcactgaccaaatgttgcagtacgcagaaggtagacctgctgataccgttaagcatgtttatggtgatacgagcaaaagtggtgacatggcgactattgtgaaatcgctttccgaagaagaagtggaagagttagctgcacactattcacagcttgcgtttgttcgtgcgaaacaggacttcgatcagtcactggctgataaagggaaaatgattcacgagaaaaattgcgacagctgtcatgtggatggaggcagcgatccgatggatgaaacctcgattctggcaggccagcaaaaaggctacttgctaaccacgctcaagcagtttcatagcggtgcacgttctgtagataagaaaatggatagcgctgttaaagggttaagtgaagacgatttggttgcactgtctgaatattacgcgagtttccaataa |
| *mtrA* | CDS (cytochrome) | *V. natriegens* ATCC 14048 | atgagagcaatcattcagcggtggttagtggtgttaatactcgggtttggggtaatttcttcggtgaactctgaagacgtgaccgagcccgaggcagaacctgaagcagcaaccttatcgagagaagagatacagaccatactggatacgaagttttctgaagggaaatactctcgtcgtggtcctgatggctgtctacgatgccatgatgacacgtctgataagcctgctacgggtattttcgataatgtacatggtaaagctgccaatattcacggaccaatgaacgataagcagtgtgaggcgtgccatggtccggcgaataaccatgaacgtaatcctcgtaaaggccaagtaagggagccaatgatcacctttggtcctaactcccctgttcctgtagagaaacaaaatagcgtgtgtttgtcgtgccaccaagatgctaaacgttctacttggcacagcagtgagcacgcttttgaaggcctatcgtgctcgagctgtcaccagttgcatcaaaaggacgatcctatgatggttgctgagttgcaagcggataaatgtacagattgccatattaaggccaaatctgacattcacaaacgcagcagtcacccggttctggatggcgtaatgacctgttcgaattgtcacaacccacaccagacaatcaacgaagcgagcctcaactggtcatcaattaatgacacctgctatgagtgtcacgcggaaaaacgcggtccaaacttatgggaacatgaacccgtcacggaagattgtacaacctgtcactctcctcacggttcagttaataaagccatgttaaacaaacgtctaccaatgctatgtcaggagtgtcaccgcgttcctcatgctaacgttgcgatgcctgaaaacgacttgaaggttcgtggtggtagctgtatgaactgccatactcaagtgcatggctctaatcacccacgcggtcagacgctaagatattaa |
| *mtrB* | CDS (beta-barrel porin) | *V. natriegens* ATCC 14048 | atgaaactatcaaaaatctgtattgctatttcccttgctttatctctacaagtacacgcagaggattattcgctgcagaatagccaagaggctgataccagccgctggaattgtactgactgtagtagcgatggtttgtggagtggtgatgtcagtgtgggcgtcggttatctcgataacgacggttcaacacgattctataactggaatccacctcttcatggcactagttcagacaacaagcacctaaatgcaagcctgaatgcggatattgaacgctatgaagatgatggcttctacaatcgcttacttgttgacgacttagggctacaacgcttcttgattcaatgggaaattggagagtatgacggttttcgtttggtaacaagttacagcgaaacgccctactactggaatcgctcatcattaagtgcctaccaagggagtgacaacgtactcacgagtggagctttgagcaaatatgatcacgaagttacccgtgataccttcaagtttgagctcaagtacacgcctaaatcaccttggcaaccctatgcatcgatgaaacatgagcgtaaagaaggcacaacgtccctctattcatcgaccataccgggttatggcagcggccccggattcattcctaaggccgtcgaccatgagacactcaatacccaagtcggtatgagctacgtagaagatctctggttggttgatgttgcctatcgcggttccttcttccgcaatgacatgtcagcgctctactatggcgatacttcagacccgtatgcgaaccaactttctaacgaacctgacaacgattttcatcagctggctgtgtccggtaattatcgcgtgaaccaacaaaccttcaatggccggattttatggtcacaagctacatctgaaggaggtttactaccatttgcgttgtccccttctaactcaaatacatttcacggtgaaaccagtacttggcaagtcagtggggattatcacaacaaactatcacgtgataccgctctgaaagtgtctgctgattactccgacaaagacgacaagtctgatagaaacgatgtggttggcgcaacgcgcgaaagctatgaccgcaccaaaacaaaactggaagcctcggtcactcatcgcctcaacagcgatatcagattgaacgcgggctatgactacacgcacgacaaacgagagtacgctgatcgtaaaagtattgatgagcaaagtttgcatttcggagctcgttatcgtccagaagcgccatggcaattaggcggaaaattatcctactcattccgagatggatcttcttggcgaaattcaagtagcgactcgcctaacttacgacaatactacttagccgataaaaaccgcatagaattacgcggagacggtagttacgacatctctgaaaacgtacaactcctcgcagaagcctggtacggcaacgatgattatgacaaacctgacattggcctgagtgagggtgaagactacggttatgatttaagtgtcaatttcaacctcgctgagggcttgaatggacatgtgttctataaccaacaaatcattcgctcagaacagcaacaagcaaattcggatgtcgttggatgggatagatacaaaacaaaactgaaagacgatgtcaccaccgttggctttggggtcagtaaagacggacttcttgaagacaagttaaccctctctctggactacagctatagtcaggcagacagtaaatcgtcaacaacggccagcgggtatcagtatccggacaatgaatacaaatcatcgcgtcttgaaggtgttgcagattatgaaatctcagacaatcaaagcgttcaattcaatgttcgttacgaagattactctgaagatgattacctcttcaataatgaagtaagtaacatgggtgacgttctgcaagactacaacggcatctatggcggcgtttattggaagtatcgcttctag |
| *mtrC* | CDS (cytochrome) | *V. natriegens* ATCC 14048 | atgaacaatataaaactacttttcatcacaacccttgccatgttaatggccgcatgtggtccaggcggtaaaaatgatcaaaatacccctccgccagatgtcagttcattcagcgtcgaaatcgaaaagccgatgcttatgtcctccgacagtggtactgaaacaaagttggtggttgactttaacgttctggacggcgcggggcgtgactacacgttcgacaccgataaagactttagaatcgcggtgttaaaagcaatgccggcacgcagtgacgttgaagatactagtgaccccgcatacgactataacgggcgaaacggcaatactttctggaaaagtttccatcactccagtgatgcctcaaataacagagcatcaatggagagtgtttgggatggcgatttagagaaaacagacgacggctatcgttacacctttgcaattacagacattttgaaagtcgccgatccgtatcctgccgattccgctgataatgacgggtacattagctgggatgaaaataagttgcaccgaatcgtgttcgcttatggcgaccaagaagtcggttttacccatgtttatgagtgggtaccaggtaccagttcagacagcactgtaacgagaaatgtcattgaagacggaacctgtgcgaattgtcatatgaacgaacccctccaccatggtccgggttatcgttcaatcgacaacaacatcgccgtttgtacgtcttgtcacaacgacagtaacccaggtgcggcacccgctcgacgaccacttgcggccgttgtacaccaatatcacggcaacgtatttaaacttggcagtgacagaaatgacctgactacgtacaagcagcctgttgatgacagtgagcagttagtcactgacatagacggtcttgtcgttgagggcagcccattcccacaagatgcgcgcaactgtacgacttgtcactcgagtgacacagcaaaagcatcggatgcaaacaactggtttgaacacccaacccaagttgcctgtgaaacctgtcacttgttccgtgatcgcggagctcatgcaaatcaagttggtacagcatggattcgagacggttcccctcaaaatacttgtaccggatgtcatcgcccttatgagcgtgacgaaaatggcgacccaatcattggcgaagacgcaagcagaagcgctaaaacggttcacatgatgcgtctggcaaacctatcaaccgccagagacacgttagacgtctcgattgaacaagcgcgtttcattgatgatagcttcgaagttcaagtacgcatcatgaaagacggtgtgggtataagctcgatggctgacattaggccttatatcaataatgacgatcacttctatcttcttcttaactgggataacggtgaaggacctctattgtcatatttaagtccgttaactttctcgtcaccatcgccgctcccaaccgttgatctgaacatgtatggagacggtgcggttggcgatggctgtgttgcagaaggtgatggtatgttcacctgtcacaaagacctcagtgcttctgatatcaaaccaaccagcacttcgaaactgactgtcaacattgcagatgtacctctatgtgctgacagaagagaaggcgtgctaagtgagtgtgtcaactttaccggcattgatctaatcaagagcccgttcatgattgcagcaaataacacgtcatcagccttcggcattggtggtgtggaaattcctcatagtcttccaatgggcgcggatattgagagctgtaacggttgtcataaagagctgacaatccacgctgactcccatgcggcgactgatttccagcaatgtaaaaattgccataactcagaacgtccatctttttacacaggtataccggcagatctgaaataccacgtgcactcgtatcatgcgtttggatctcatcgtaatggtgaggcggtattcccaggagcaattaacaattgtgaatcttgccatacatcgtctcaatttaacctgccaagtcaggaaaatacgcgcccatcattggcgatgaatacagcaggtgaaacaaaatacttcagcccggctctagtggcctgtggtgcatgtcaccttgaatcagcgctcgctgacgcagatccagaaactgtagctggtgatgcaatactgagccatatgataaataacggcgcagtctttggtgctgacacagcagcagaagcgatgggttctgagcaatgttcgacatgtcacgctatcgggcagtctcagggcgtagataaggttcacaaggtatatgactaccgttga |

Supplemental Table 5: List of genetic parts used in chromosomal insertions. The different compositions of these parts that were used to assemble gene expression cassettes are detailed in **Supplemental Table 4**.

**Supplemental Table 6**

| **Primer** | **Sequence** |
| --- | --- |
| MC335 | agcctttacataggtctgacagtc |
| MC336 | tcatcatggcatcgtagacagc |
| MC339 | cttccacttcttcttcggaaagc |
| MC371 | ggttctcaacttctccctcttttactag |
| MC372 | ccgatgagttggtttgtgcg |
| MC373 | gaccgcttttacagagtcagc |
| MC389 | caaagttggtggttgactttaacg |
| MC390 | ggtctgaatattgtggcgatgatc |
| WC27 | cgattggtatcacctcataccctgac |
| WC28 | ggtaaagccagtcacttcacaacc |
| WC68 | cgttcgccaaactgtgcac |
| WC69 | gaccaagtaccgtttggacgtc |
| WC93 | cgcgggtcaacttcaatgctg |
| WC94 | cgcgactgggtgttagagc |

Supplemental Table 6: Primer sequences

**References**

1.	Park Y, Espah Borujeni A, Gorochowski TE, Shin J, Voigt CA. 2020. Precision design of stable genetic circuits carried in highly-insulated E. coli genomic landing pads. Mol Syst Biol 16:e9584.

2.	Meyer AJ, Segall-Shapiro TH, Glassey E, Zhang J, Voigt CA. 2019. Escherichia coli “Marionette” strains with 12 highly optimized small-molecule sensors. Nat Chem Biol 15:196–204.

3.	Bhakta SP. 2024. A Bacterial Toolkit for Rapid Prototyping of Multicistronic Genetic Circuits from Interchangeable Parts. Rice University.

4.	Sun Y, Xu J, Zhou H, Zhang H, Wu J, Yang L. 2023. Recombinant Protein Expression Chassis Library of Vibrio natriegens by Fine-Tuning the Expression of T7 RNA Polymerase. ACS Synth Biol 12:555–564.

5.	Wu F, Chen W, Peng Y, Tu R, Lin Y, Xing J, Wang Q. 2020. Design and Reconstruction of Regulatory Parts for Fast-Growing Vibrio natriegens Synthetic Biology. ACS Synth Biol 9:2399–2409.
